# A High-Sensitivity Stopped-Flow EPR System to Monitor Millisecond Conformational Kinetics in Spin-Labeled Proteins

**DOI:** 10.1101/2025.03.26.645583

**Authors:** Alexander M. Garces, Richard R. Mett, Candice S. Klug, Jason W. Sidabras, Michael T. Lerch

## Abstract

Electron paramagnetic resonance (EPR) spectroscopy is a powerful tool for studying biological systems, with applications in drug discovery, protein dynamics, membrane biology, and enzyme mechanisms. However, sample volume requirements and sensitivity limitations have historically constrained time-resolved measurements of protein dynamics using stopped-flow (SF) EPR spectroscopy. To address these challenges, we developed a high-sensitivity SF EPR system featuring a custom dielectric resonator, an optimized low-volume sample tube geometry design, and the SF mixer assembly integrated into the resonator housing. This system significantly reduces sample requirements for the investigation of protein conformational dynamics on the millisecond timescale. We demonstrate its capabilities through two applications: the analysis of T4 lysozyme unfolding kinetics, which revealed site-specific variations in the folding pathway, and the measurement of ligand-induced conformational changes in the β2 adrenergic receptor, a challenging membrane-protein system. This advancement broadens the applicability of SF EPR to complex, biomedically relevant proteins, facilitating studies of protein-protein and protein-ligand interactions in diverse biological processes.

## 1. Introduction

Proteins are intrinsically dynamic molecules, and the role of protein motion in biological function is well-established in enzyme catalysis, membrane transport, protein-protein or protein-nucleic acid interactions, and signal transduction (Borgia et al., 2018; Janetzko et al., 2022; Manglik & Kobilka, 2014; Nam & Wolf-Watz, 2023; Shcherbakov et al., 2023; Zeymer et al., 2016). Equilibrium fluctuations in protein structure take place over a wide range of time scales and range in amplitude from subtle changes in side chain rotamers to collective motions of entire domains. Functional motions such as changes in loop position, rigid body motion of helices, and local secondary structure generally occur on the microsecond timescale or slower. These conformational fluctuations are described by an energy landscape that defines the relative probabilities of the states that comprise the conformational ensemble as well as their interconversion paths and kinetics (Henzler-Wildman & Kern, 2007; Mishra & Jha, 2022). A complete description of the molecular mechanisms of protein function encompasses defining the features of the conformational energy landscape, including both the relative populations and lifetimes of states, and how the landscape is modulated through binding interactions or changes in cellular conditions. The time dependence of biomolecular structural changes remains underexplored and is required to define the role of transient states and dynamically driven allostery in protein function (Astore et al., 2024).

To elucidate the energy landscape for proteins in solution, site-directed spin labeling (SDSL) electron paramagnetic resonance (EPR) is a particularly attractive option due to its high sensitivity, an intrinsic timescale that is favorable for measuring protein dynamics, and no inherent limitations on the size or complexity of the protein system that may be studied. Importantly, SDSL EPR can be applied under physiologically relevant conditions, making it a versatile tool for studying protein dynamics. SDSL-EPR is an established approach to site-selectively map secondary and tertiary structure, identify flexible regions, determine oligomerization and protein complex formation, and monitor protein dynamics (Altenbach et al., 2015; Bordignon & Bleicken, 2018; Cafiso, 2014; Claxton et al., 2015; Galazzo & Bordignon, 2023; García-Rubio, 2020; Goldfarb, 2022; Jeschke, 2018; Roser et al., 2016; Torricella et al., 2021). The continuous wave (CW) EPR spectrum of a spin-labeled protein is sensitive to motion on the 0.1–100 ns timescale and encodes information on local backbone dynamics, structure, and topology. Conformational exchange in the microsecond-to-millisecond range is slow on the EPR timescale, and each conformation gives rise to a distinct spectral component that provides information on the local tertiary fold and backbone dynamics. The techniques of pulsed saturation recovery (Bridges et al., 2017) and saturation transfer EPR (Chen et al., 2023; Hustedt & Beth, 2004; James et al., 2012) can access exchange rates on this timescale through the saturation behavior of the electron spin-lattice relaxation time. Pulsed saturation recovery can measure exchange rates on ∼1– 100 µs timescale. In cases where internal motion of the spin label and rotational motion of the protein can be sufficiently suppressed, saturation transfer EPR extends the accessible time window to milliseconds. In most cases, these techniques have a practical upper limit for measuring conformational exchange kinetics of ∼1 ms. Rapid-mixing methods extend the time domain for detection by SDSL EPR into the ms– s timescale, enabling exploration of large (multi-Ångstrom) collective motions of proteins such as domain movements and rigid-body motions of helices. Stopped-flow (SF) EPR spectroscopy monitors changes in EPR signal, in real time, following rapid mixing of reactants and has demonstrated substantial utility across numerous applications, including studies of metalloprotein activation and free radical biology (Bennett, 1990; Gilbert et al., 1997; Kuwabara et al., 2018; Lisovskaya et al., 2021; Pötsch et al., 1995; Sen et al., 2006) as well as the kinetics of conformational changes in spin-labeled proteins and nucleic acids (DeWeerd et al., 2001; Grigoryants et al., 2000, 2004; Qu et al., 1997; Shin et al., 1993). SF EPR combined with SDSL enables time-resolved monitoring of site-specific protein conformational changes triggered by ligand binding, membrane interactions, or protein-protein interactions, or changes in solution conditions (e.g., pH). Despite the demonstrated ability to provide unique insights into functional dynamics and protein folding pathways (Qu et al., 1997; Shin et al., 1993), applications of SF EPR on membrane proteins and protein complexes have been limited by the significant amount of spin-labeled protein required for data collection.

In this study, we describe an SF EPR system capable of resolving millisecond timescale conformational exchange rates in spin-labeled proteins. To our knowledge, this system achieves the lowest sample consumption per shot reported for a SF EPR system, while maintaining high detection sensitivity through implementation of a custom resonator and novel sample tube geometry. Two initial applications are presented to showcase this advancement: (i) site-specific folding kinetics of the well-characterized model protein T4 lysozyme (T4L) in urea, and (ii) the rate of agonist binding and activation of the β2 adrenergic receptor (β2AR). The results highlight the exceptional sensitivity of the novel SF EPR system, enabling both analysis of T4L folding pathways through site-specific kinetic measurements and the detection of time-resolved allosteric conformational changes in a complex membrane protein system.

## 2. Results

The integrated SF EPR system comprises several custom components designed to minimize sample requirements through reducing sample volume consumption and maximizing signal sensitivity. The key components are a custom resonator housing and an SF mixer assembly, as well as a novel dielectric X-band resonator and sample tube.

### 2.1 SF EPR system design

This SF EPR system incorporates a µSFM instrument, (BioLogic; Seyssinet-pariset, Rhone-Alpes, France) selected for its superior accuracy in low-volume applications. Although this driver system was previously utilized in a rapid-freeze quench system (Pievo et al., 2013), it had not been integrated into SF EPR applications prior to this work. The system employs a micro-Berger-Ball mixer capable of efficiently mixing sample volumes as small as 3 µL (Berger et al., 1968; Kathuria et al., 2013; Lassmann et al., 2005). The driver system included with the commercial µSFM instrument is a dual syringe design that utilizes independent stepper motors for each syringe, allowing for mixing ratios of 1:1 to 1:9. A hard-stop valve is incorporated after the resonator to terminate sample flow after each shot. System operation is managed through Bio-Kine software (BioLogic; Seyssinet-pariset, Rhone-Alpes, France), which communicates with the Elexsys E500 spectrometer (Bruker; Billerica, Massachusetts) via a 5-volt transistor-transistor logic pulse for precise data acquisition timing.

To optimize this system for SF EPR, two critical parameters that govern SF system performance were considered: dead volume and dead time. Dead volume encompasses the space between the mixer outlet and the center of the EPR-active volume, while dead time represents the time required for the sample to travel from the mixer end to the center of the detection point. Optimization of these parameters is crucial for maximizing time resolution and minimizing sample consumption.

Through strategic design choices, we achieved substantial improvements in system performance. A key optimization involved the careful positioning of the driver system horizontally on a cart, allowing partial insertion into the spectrometer’s electromagnet. This arrangement isolated the driver system motors from the spectrometer’s electromagnetic field while minimizing the required priming volume. To implement this configuration, we developed a custom driver system adaptor and mixer adaptor, connected via PTFE (1.6 OD × 0.5 mm ID) tubing approximately 40 mm in length. This design reduced the priming volume to approximately 30 μL per syringe.

The mixer adaptor is integrated directly into the resonator housing assembly (Fig. 1), yielding a geometrically calculated dead volume of 6.93 µL and a dead time of approximately 3.8 ms at the fastest flow rate available to the biologic driver system of 1.8 mL/s. The sample volume in the active region of the resonator is 4.39 μL using the sample tube described below. The age distribution width—defined as the time required to travel through the active region—is 2.4 ms at a flow rate of 1.8 mL/s. Lastly, a custom adaptor was fabricated to attach PTFE (1.6 mm OD × 0.5 mm ID) tubing that exits the resonator, to the hard-stop valve (BioLogic; Seyssinet-pariset, Rhone-Alpes, France). The custom adapters were designed using Autodesk® Inventor (San Francisco, California) and manufactured via stereolithography using a FormLabs (Somerville, Massachusetts) Form 3B printer. Temperature control is maintained through a RHC-400 Circulating Bath, (Cole-Parmer;Vernon Hills, Illinois) connected to the μSFM system’s integrated cooling channels surrounding the syringes.

**Figure 1.**
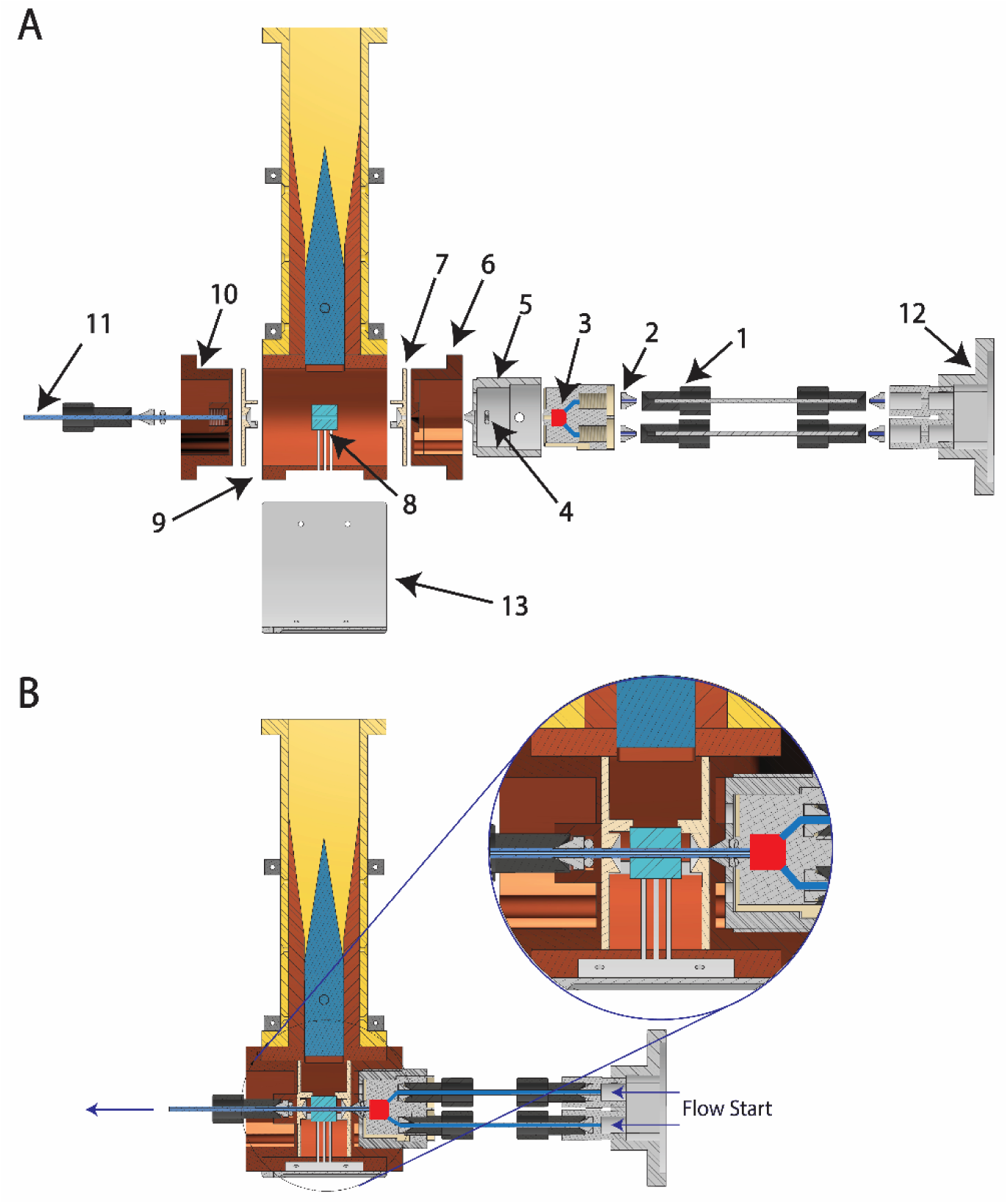
SF EPR assembly. (**A**) Expanded and (**B**) assembled resonator and mixer assembly. Individual components are (**1**) FPLC fittings, ¼-28 thread, 1/8’’ ID; (**2**) FPLC ETFE ferrule; (**3**) micro-Berger-Ball mixer; (**4**) silicone O-ring; (**5**) mixer adaptor; (**6**) top copper resonator cap; (**7**) Rexolite spacer for the sapphire dielectric; (**8**) sapphire dielectric resonator; (**9**) cylindrical copper housing; (**10**) bottom copper resonator cap; (**11**) PTFE tubing, 0.5 mm ID, 3.18 mm OD; (**12**) driver system adaptor (connects directly to μSFM outlet); (**13**) modulation coil housing.

### 2.2 Resonator design

A dielectric resonator operating at X-band was designed to maximize the signal intensity of low-volume, low-concentration samples (Fig. 1). Simulations using Ansys High Frequency Structure Simulator (Ansys Electronics Desktop 2024R1; Canonsburg, Pennsylvania) were performed to design and optimize the sapphire dielectric resonator geometry, the oversized shield, and the custom sample tube geometries. The core component of the dielectric resonator is a grown sapphire crystal, which was precisely machined and polished by Insaco (Quakertown, PA) to form a hollow cylindrical structure. The high-sensitivity resonator was assembled in-house following the design principles of Hyde et al. (Mett & Hyde, 2022). Mainly, that the dielectric resonator has the highest EPR sensitivity when the effects of the shield are negligible on the dielectric resonator frequencies and, in turn, ohmic losses. This design principle leads to an oversized—compared with typical dielectric resonator shields—metal cavity that acts as both a coupling mechanism and a shield. The oversized resonator shield was crafted from solid tellurium copper, machined into a hollow cylinder with three modulation slots integrated into its body. The geometry design ensured that the sapphire resonator and iris perturbation do not excite resonant modes within the cavity shield.

The resonator is attached to a WR-90 waveguide with an oversized iris (12.65 × 12.65 mm^2^) that reduces reactive coupling (low capacitive and inductive coupling). Match variation is accomplished through a Gordon coupler equipped with a Rexolite plug (Mett & Hyde, 2022). One advantage of a Gordon coupler is that when in conjunction with an oversized iris, it is a purely resistive coupling system. This means that moving the Gordon coupler away from the iris changes only the magnitude of the coupling, resulting in no frequency shifts while coupling (Mett & Hyde, 2022). Eliminating frequency shifts during coupling is especially important for long-term experiments and multi-shot signal averaging such as is needed for SF EPR.

A custom sample tube based on the AquaStar geometry previously developed for higher volumes (Sidabras et al., 2017) was fabricated using new ultra-precision 3D printing techniques. This reduced AquaStar geometry cross section, illustrated in Figure 2, was manufactured using a Boston Microfabrication (Maynard, Massachusetts) S140 3D resin printer, which employs projection micro stereolithography technology to achieve an exceptional printing resolution of 10 µm. The precise manufacturing capabilities of the projection micro stereolithography technology ensure consistent quality and dimensional accuracy, which are crucial for reliable spectroscopic measurements. The sample cell design improves the EPR signal intensity by reducing electric field losses within the aqueous sample. The sample tube was printed using HTL (Boston Microfabrication; Maynard, Massachusetts), a high-performance photopolymer engineering material.

**Figure 2.**
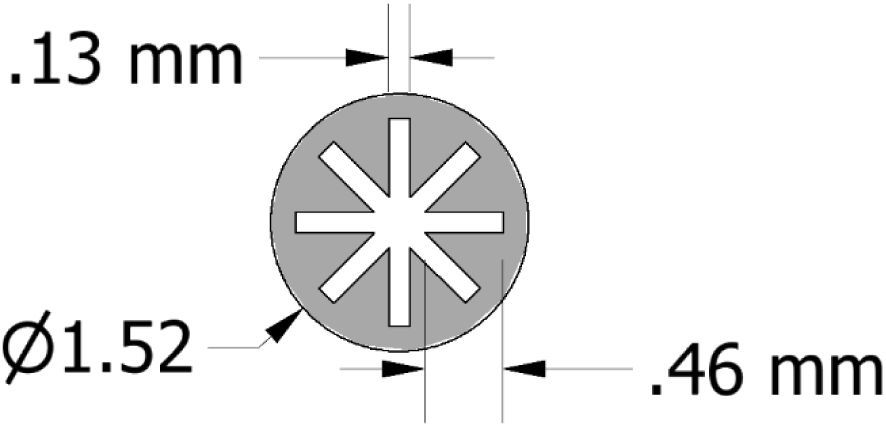
Cross-section schematic of the AquaStar sample tube.

The positioning of the sapphire crystal within the copper housing is maintained by two Rexolite spacers, which are secured by copper caps screwed into the main housing (Fig. 1). This sapphire dielectric resonator configuration is crucial to the SF EPR system, as it provides an optimal geometric compromise between achieving maximum signal intensity and allowing the Berger-Ball mixer to be positioned as close as possible to the active region of the resonator.

For modulation capabilities, custom modulation coils were hand-wrapped around both sides of the resonator. These coils were wrapped around a custom designed holder, which was created using AutoCAD Inventor (San Francisco, California) and produced on a Bambu Lab (Shenzhen, China) X1 Carbon 3D printer. These custom-made coils provide modulation capabilities of up to 3 Gauss, which is sufficient for CW SF studies.

### 2.3 SF EPR system performance

To evaluate the performance of the novel integrated low-volume resonator and stopped flow system, the reduction of TEMPOL (4-hydroxy-2,2,6,6-tetramethyl-piperidine-N-oxyl) by sodium dithionite was monitored. This reaction is commonly used as a benchmark for the characterization of SF EPR system performance, including dead time determination (Grigoryants et al., 2000; Lassmann et al., 2005; Schubert et al., 2016). The reaction kinetics were monitored at a fixed magnetic field position corresponding to the low-field line maximum of the TEMPOL spectrum (Fig. 3A). SF EPR experiments reveal a characteristic signal profile composed of several distinct phases (Fig. 3B). Each experiment begins with a programmed delay period wherein data are collected before the driver system initiates flow of the individual reagents. The flow sequence starts with the opening of the hard stop valve followed by synchronized activation of the two syringe drivers. As the mixed sample begins to flow through the resonator, a sharp signal increase is observed as TEMPOL enters the resonator’s active region. This is followed by a stable plateau phase, wherein steady-state equilibrium is established between the rate of sample replacement in the active region and the ongoing reaction kinetics (i.e., continuous-flow condition). Finally, when the predetermined shot volume is reached, the flow terminates and the hard-stop valve closes, marking the beginning of the reaction decay phase. The complete signal trace, encompassing all phases from initial flow to final decay, is recorded by the spectrometer’s data acquisition system for subsequent analysis (Fig. 3B). This experimental sequence captures both the mixing dynamics and reaction kinetics in a single measurement, providing a comprehensive view of the reaction progress.

**Figure 3.**
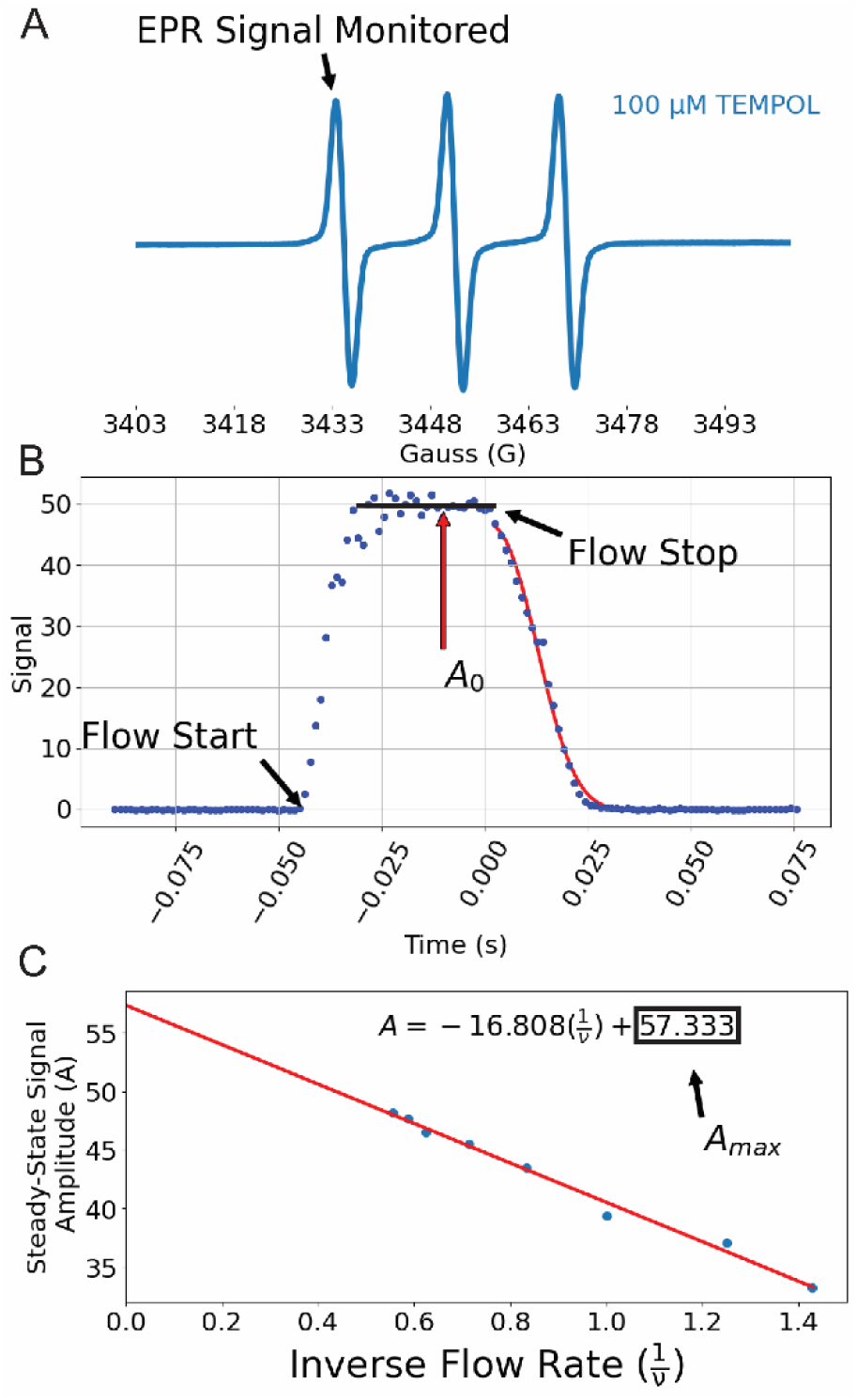
Experimental estimation of SF EPR dead time using the reduction of TEMPOL by sodium dithionite. (**A**) X-band EPR spectrum of 100μM TEMPOL. The black arrow indicates the low field line monitored in subsequent SF EPR experiments. (**B**) Complete SF EPR trace of 1 mM TEMPOL reduced by 50 mM sodium dithionite at a flow rate of 1.8 mL/s. A stretched exponential fit to the signal decay after flow termination is shown in red. (**C**) The average steady state signal amplitude (*A*) is plotted versus the inverse flow rate. A fit to a linear equation (inset) extrapolated to 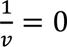 yields the theoretical signal amplitude at infinite flow rate (*A*_*max*_).

SF EPR measurements of the reduction of TEMPOL by sodium dithionite were conducted using flow rates ranging from 0.7 mL/s to 1.8 mL/s (Fig. S1). Performing this reaction at different flow rates results in different aging times for the mixed sample in the active region of the resonator and therefore different steady-state signal amplitudes (*A*). A plot of *A* as a function of inverse flow rate 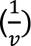 is well-fit to a linear equation, yielding an extrapolated signal amplitude at an infinite flow rate (*A*_*max*_) of 57.33 (Fig. 3C). Using the parameters *A*_*max*_, the amplitude *A* at the maximum flow rate, and the reaction decay time constant τ, an estimated dead time of 3.46 ms was determined (see Materials and Methods). This experimental value is in remarkably close agreement with the geometrically calculated dead time of 3.8 ms, confirming the ability of the system to monitor reactions on the millisecond timescale.

There has been some discrepancy in the treatment of the kinetics of the reaction between TEMPOL and sodium dithionite (Grigoryants et al., 2000; Lassmann et al., 2005; Schubert et al., 2016), and although previous studies under comparable conditions utilized zero- or first-order models to fit the relaxation profiles, the reported fits are imperfect at similar TEMPOL and sodium dithionite concentrations (Grigoryants et al., 2000; Lassmann et al., 2005). We systematically evaluated three fitting models to fit our data: linear (zero-order), single exponential (pseudo-first order), and compressed exponential. Analysis of residuals revealed systematic curvature for both linear and exponential fits, indicating inadequate modeling of the reaction kinetics. To reconcile the imperfect fits of zero- and first-order models to the TEMPOL-dithionite reaction kinetics (Fig. S3), we used a hypothesis-driven approach employing a compressed exponential function (*equation 1*) (Lukichev, 2019). The compressed exponential model provided superior fit quality, reflecting its ability to account for spatiotemporal heterogeneity present in the active region of the resonator.

### 2.4 T4L folding kinetics determined with SF EPR

To demonstrate the capacity of SF EPR spectroscopy to provide site-specific conformational exchange rates with high temporal resolution, we investigated the local folding kinetics of T4L in urea at three distinct regions of the protein. T4L has served as a useful model system to illustrate the role of intermediates and subdomain cooperativity in protein-kinetic folding pathways (Cellitti, Bernstein, et al., 2007; Cellitti, Llinas, et al., 2007; Llinás & Marqusee, 1998; Manuel Llinás et al., 1999; Parker & Marqusee, 1999), and has been used extensively to evaluate novel EPR technology (Bridges et al., 2010; Chen et al., 2023; Georgieva et al., 2012; Grosskopf et al., 2024; López et al., 2014; Mccoy & Hubbell, 2011). Previous studies have characterized the equilibrium and kinetic unfolding pathway of T4L using chemical denaturants such as urea to modulate the folding-unfolding equilibrium (Cellitti, Bernstein, et al., 2007; Cellitti, Llinas, et al., 2007; Llinás & Marqusee, 1998; Parker & Marqusee, 1999, 2001). The equilibrium urea denaturation of T4L determined with circular dichroism follows a two-state model with a midpoint at ∼5.5 M urea; nearly complete loss of secondary structure and tertiary fold occurs at >7 M urea (Mccoy & Hubbell, 2011). In contrast, investigation of T4L folding and unfolding pathways through native state hydrogen exchange and SF circular dichroism were consistent with a four-state model featuring an intermediate state in which only the C-terminal domain maintains its structure (Cellitti, Bernstein, et al., 2007; Cellitti, Llinas, et al., 2007; Llinás & Marqusee, 1998).

We monitored protein folding kinetics with the spin label R1 attached at three distinct sites: the H helix (V131R1), C helix (N68R1), and B helix (S44R1) (Fig. 4A). At solvent exposed sites such as these, the spin label side chain R1 typically has weak or no interactions with the protein and introduces little structural perturbation (Fleissner et al., n.d., 2009; Guo et al., 2008; Mchaourab et al., 1996). Thus, R1 serves as a sensor of local structure at these residues. CW spectra reflect the rotational correlation time of the nitroxide on the 0.1–100 ns timescale, which includes contributions from internal side chain motions, local protein fluctuations, and—for small, soluble proteins such as T4L— overall protein rotational diffusion (López et al., 2009, 2012). Here, experiments were performed in the absence of a viscogen, and the similarity of the CW spectra for S44R1, N68R1, and V131R1 in 0 M urea (Fig. S2) indicates that rotational motion of the protein is the predominant contribution to the lineshape (Fig. 4B). Pronounced narrowing of the three resonance lines is observed for all three constructs in response to urea-induced unfolding. These spectral changes reflect an increase in nitroxide mobility on the ns timescale consistent with unfolding local to each of the spin-labeled sites.

**Figure 4.**
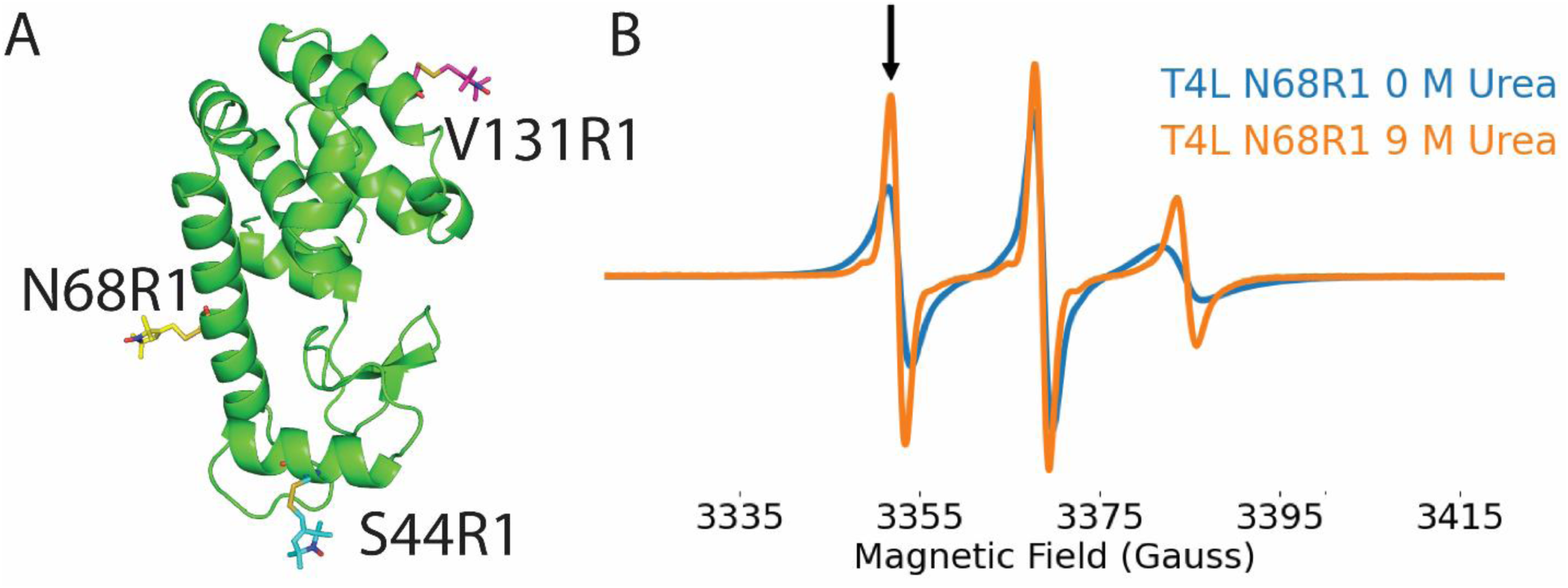
Urea dependence of T4 lysozyme CW-EPR spectra. (**A**) Crystal structure of T4L (PDB: 1L60) with the R1 side chain shown at residues 44, 68, and 131. (**B**) CW EPR spectra of T4L N68R1 in 0 M and 9 M urea. The black arrow indicates the field position monitored in SF EPR experiments.

Examples of SF EPR relaxation profiles for T4L are shown in Fig. 5A. The signal amplitude at the maximum of the low field line (black arrow, Fig. 4B) was monitored in all T4L SF EPR experiments. A decrease in the low field line signal amplitude indicates a shift toward the natively folded (N) state. Site-specific kinetics are readily apparent in the relaxation profiles collected for the different constructs at the same urea concentration (e.g., Fig. 5A). The mean and 95% confidence interval of the relaxation rates for each T4L construct at different urea concentrations are shown in Supplemental Tables S4–S6. Although most relaxation profiles were consistent with a single exponential process, some required a bi-exponential model to generate an adequate fit as described in the Materials and Methods section (e.g., Fig. 5B). The natural logarithm of the rate constant (k) is plotted as a function of urea concentration for each construct in Fig. 6. These plots display the expected chevron-like behavior, with turnover in both the folding and unfolding limb as previously reported for T4L (Cellitti, Bernstein, et al., 2007). In order to adequately fit the turnover in both limbs, Marqusee et al. (Cellitti, Bernstein, et al., 2007) present a four-state sequential folding pathway that includes both a folding and unfolding intermediate. The transition between these intermediate states is the rate-limiting step in the folding and unfolding process. A fit to this four-state model for the folding pathway (Equations 5–11) yielded equilibrium constants for the U ↔ I and N ↔ J equilibria and microscopic rate constants for the I → J and J → I transitions, provided in Table 1. Notably, high correlations between parameters lead to broad confidence intervals for some parameters. Together, these data highlight the ability of the SF EPR system to site-specifically monitor complex reaction pathways by capturing time-dependent changes in conformational equilibria across nearly 200 relaxation profiles, all while requiring only ∼100 nmol of protein for the entire dataset.

**Figure 5.**
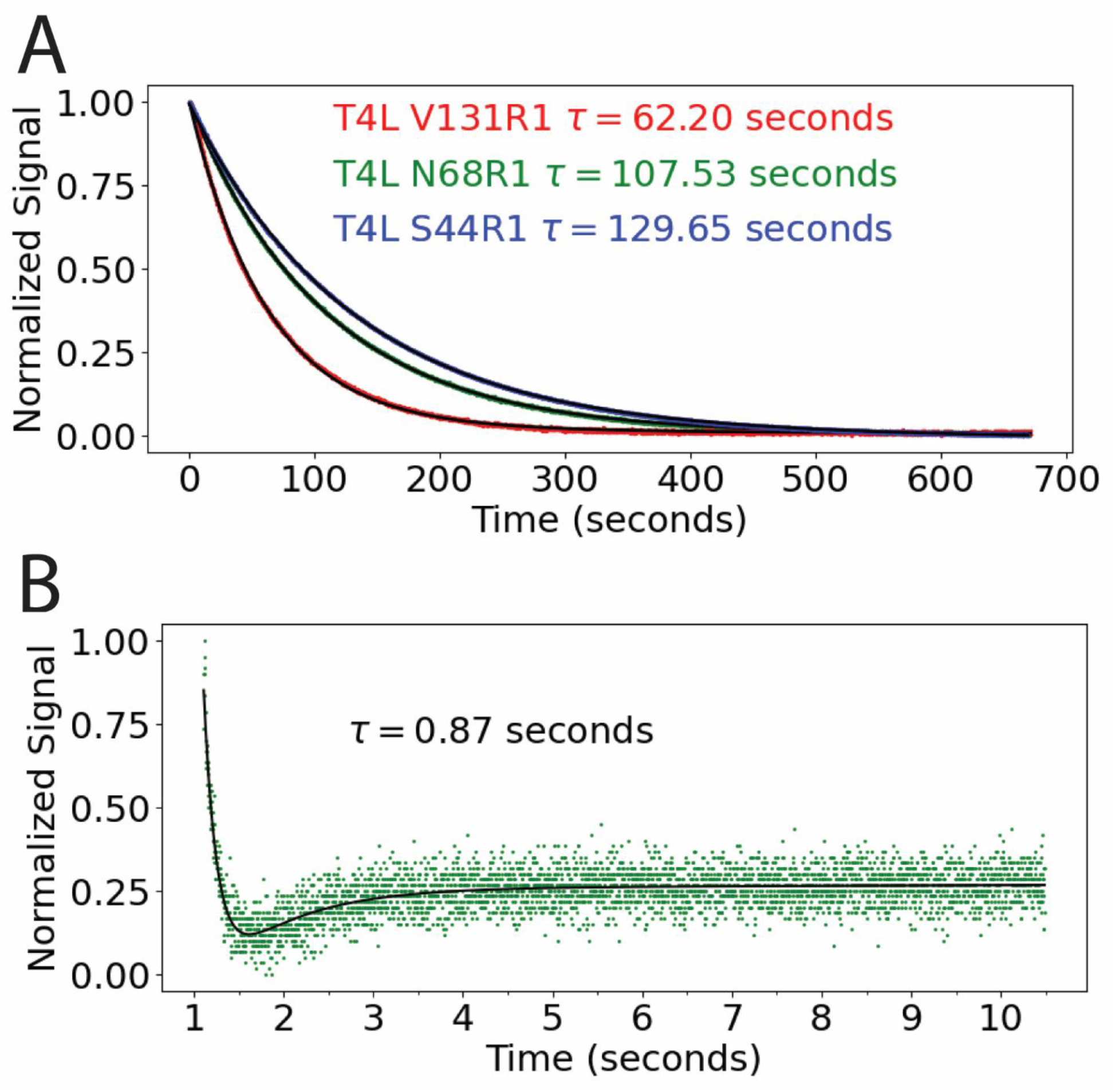
Examples of SF EPR relaxation profiles for T4L. (**A**) SF EPR relaxation profiles of T4L S44R1 (blue), N68R1 (green), and V131R1 (red) folding at 5 M urea fitted to an exponential decay (black). (**B**) The SF EPR relaxation profile of T4L N68R1 at 3 M urea illustrates the bi-phasic behavior observed in a subset of experiments. This relaxation profile is well-fit to a bi-exponential decay (black).

**Figure 6.**
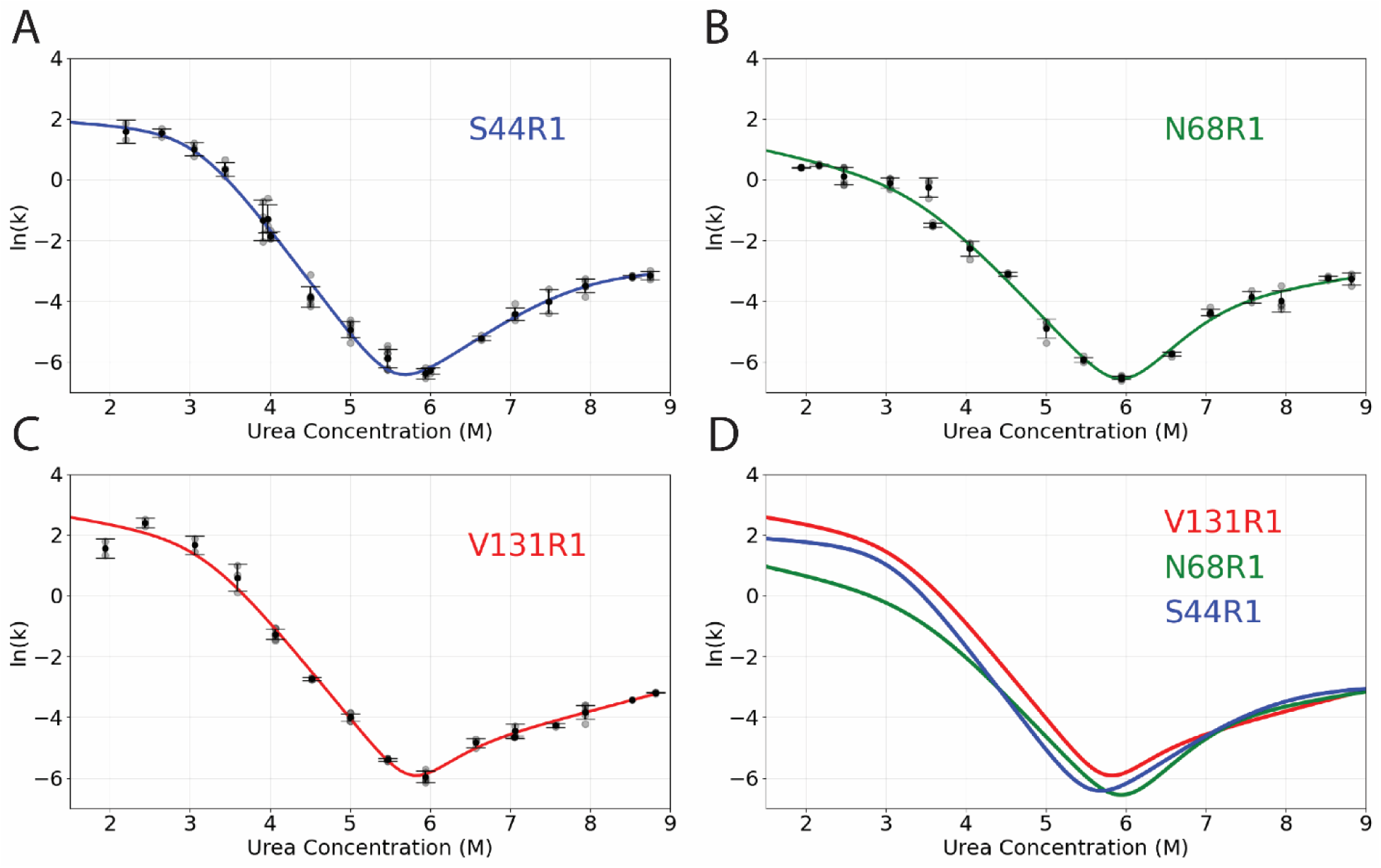
Urea-dependent kinetics of T4L folding at three distinct regions of the protein: helix H (V131R1), helix C (N68R1), and helix B (S44R1). The natural logarithm of the relaxation time constants determined from fits to individual SF EPR relaxation profiles (grey dots) are plotted versus urea concentration for (**A**) S44R1, (**B**) N68R1, and (**C**) V131R1; at each concentration, means are shown as black dots and error bars indicate standard deviation of ln(k) values at each urea concentration. Fits to a four-state model (color-coded as indicated) yielded kinetic and thermodynamic parameters characterizing the folding reaction pathway. Fitted parameter values are listed in Table 1. (**D**) An overlay of the fits for each construct illustrates the site specificity of the kinetics.

**Table 1:**
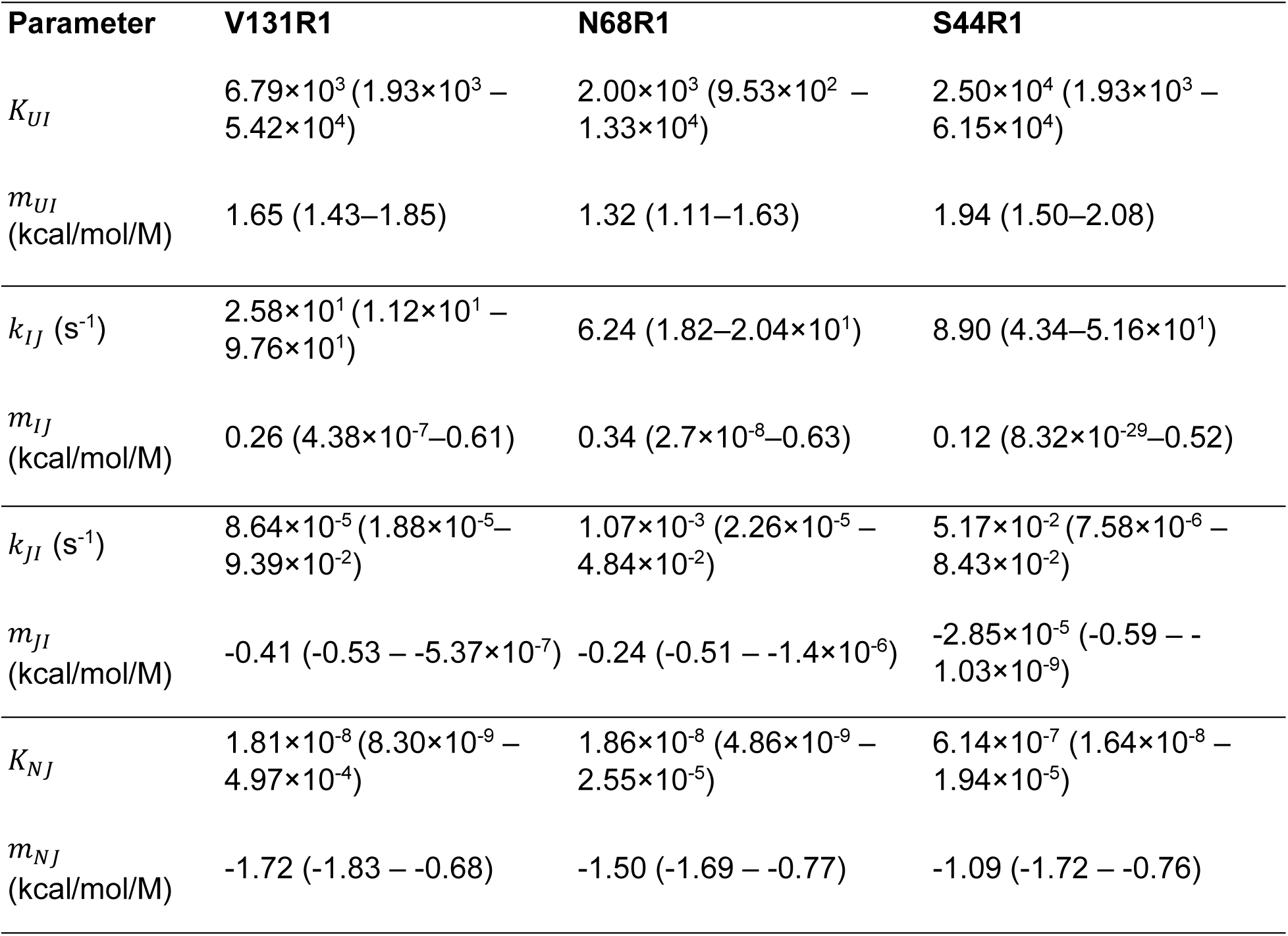
Fitted Parameter Values From Analysis of T4L Folding Kinetics (Figure 6) According to a Four-State Model.

| Parameter | V131R1 | N68R1 | S44R1 |
| --- | --- | --- | --- |
| $K_{UI}$ | $6.79 \times 10^3 (1.93 \times 10^3 - 5.42 \times 10^4)$ | $2.00 \times 10^3 (9.53 \times 10^2 - 1.33 \times 10^4)$ | $2.50 \times 10^4 (1.93 \times 10^3 - 6.15 \times 10^4)$ |
| $m_{UI}$<br>(kcal/mol/M) | 1.65 (1.43–1.85) | 1.32 (1.11–1.63) | 1.94 (1.50–2.08) |
| $k_{IJ}$ (s<sup>-1</sup>) | $2.58 \times 10^1 (1.12 \times 10^1 - 9.76 \times 10^1)$ | 6.24 (1.82–2.04×10<sup>1</sup>) | 8.90 (4.34–5.16×10<sup>1</sup>) |
| $m_{IJ}$<br>(kcal/mol/M) | 0.26 (4.38×10<sup>-7</sup>–0.61) | 0.34 (2.7×10<sup>-8</sup>–0.63) | 0.12 (8.32×10<sup>-29</sup>–0.52) |
| $k_{JI}$ (s<sup>-1</sup>) | $8.64 \times 10^{-5} (1.88 \times 10^{-5} - 9.39 \times 10^{-2})$ | $1.07 \times 10^{-3} (2.26 \times 10^{-5} - 4.84 \times 10^{-2})$ | $5.17 \times 10^{-2} (7.58 \times 10^{-6} - 8.43 \times 10^{-2})$ |
| $m_{JI}$<br>(kcal/mol/M) | -0.41 (-0.53 – -5.37×10<sup>-7</sup>) | -0.24 (-0.51 – -1.4×10<sup>-6</sup>) | -2.85×10<sup>-5</sup> (-0.59 – -1.03×10<sup>-9</sup>) |
| $K_{NJ}$ | $1.81 \times 10^{-8} (8.30 \times 10^{-9} - 4.97 \times 10^{-4})$ | $1.86 \times 10^{-8} (4.86 \times 10^{-9} - 2.55 \times 10^{-5})$ | $6.14 \times 10^{-7} (1.64 \times 10^{-8} - 1.94 \times 10^{-5})$ |
| $m_{NJ}$<br>(kcal/mol/M) | -1.72 (-1.83 – -0.68) | -1.50 (-1.69 – -0.77) | -1.09 (-1.72 – -0.76) |

### 2.6 Ligand-induced activation of the β2AR

To further demonstrate the utility of this system, conformational exchange rates were measured for the β2-adrenergic receptor (β2AR), a biomedically important G-protein coupled receptor (GPCR). The outward tilt of transmembrane helix 6 (TM6) is the largest conformational rearrangement associated with β2AR activation and is required for this receptor to productively couple to its cognate signal transduction partners (transducers). The effects of ligand- and transducer-binding on TM6 conformation have been delineated structurally and spectroscopically (Gregorio et al., 2017; Manglik et al., 2015; Rasmussen, Choi, et al., 2011; Rasmussen, Devree, et al., 2011a; Rasmussen et al., 2007). In brief, activating ligands and transducers stabilize an outward-tilted conformation of TM6, whereas inactivating ligands stabilize an inward conformation of TM6.

Here, R1 is placed at the cytoplasmic end of TM6 (C265R1) to monitor the equilibrium between inward and outward conformations of the helix. CW spectra of β2AR C265R1 were collected in the unliganded state and bound to the strong agonist BI-167107 (Fig. 7a), which has been shown spectroscopically and structurally to populate the outward conformation of TM6 (Gregorio et al., 2017; Ma et al., 2020; Manglik et al., 2015; Nygaard et al., 2013; Rasmussen, Devree, et al., 2011b). The spectrum of the unliganded receptor is dominated by a component reflecting immobilization of the spin label, with a second component reflecting a more mobile state. The immobile component is consistent with an inward position of TM6, in which contacts between the R1 side chain and the local protein environment would lead to reduced motion of the nitroxide. Binding of the agonist increases the relative population of the mobile component. Based on prior structural evidence, we interpret this spectral change as reflecting a shift in the TM6 conformational equilibrium toward the outward, active state, thereby relieving local contacts and resulting in an increase in nitroxide mobility.

**Figure 7.**
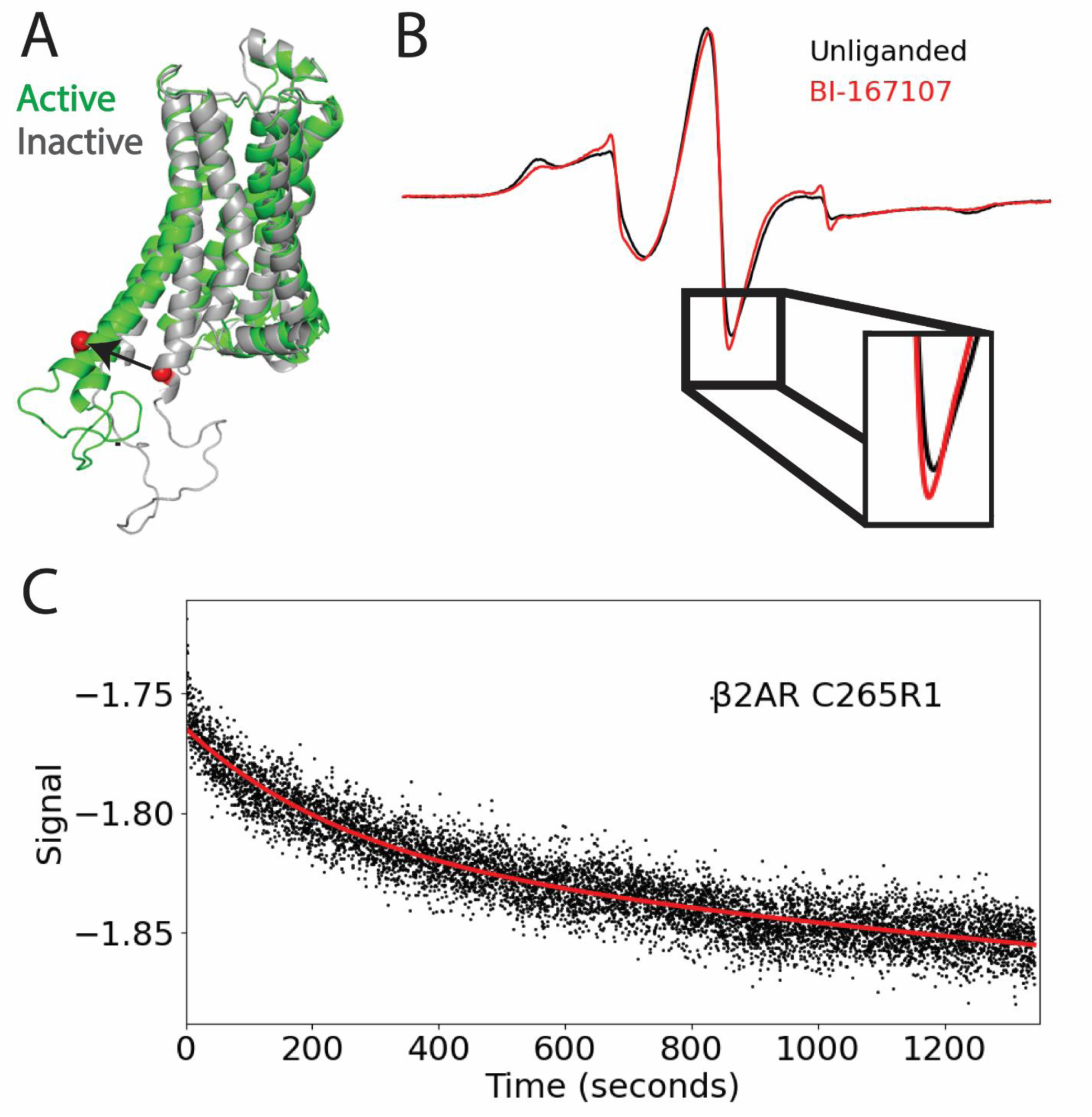
Spin-labeled β2AR C265R1, CW, and SF EPR data. (**A**) PDB: 3SN6 (active) and 5JGH (inactive), with red spheres indicating spin-labeled residue C265R1. (**B**) CW EPR spectra of β2AR C265R1 apo and β2AR C265R1 bound to BI-167107. (**C**) SF EPR β2AR C265R1 apo mixed with buffer + 1x BI-167107. 1:1 mixing ratio, with a flow rate of 1.8 mL/s.

The kinetics of TM6 conformational exchange was followed upon binding of BI-167107 to unliganded β2AR using the signal amplitude at the minimum of the center line (Fig. 7). The resulting relaxation profile was fit using a single exponential model, yielding a fitted relaxation time constant of 259 s. These data illustrate the capacity for real-time measurement of allosteric conformational changes in GPCRs and establish a foundation for exploring the role of dynamics of GPCR activation.

## 3. Discussion

### 3.1 Performance of the system

This study presents a novel X-band dielectric resonator tailored to minimize sample requirements for measuring reaction kinetics of spin-labeled proteins in the milliseconds-to-seconds regime. The custom resonator assembly includes a sapphire resonator, an oversized cavity shield, and a custom 3D printed low-volume sample tubes with AquaStar geometry. Four principles highlight the novel features of this low-volume resonator: (i) use of a single crystal sapphire reduces background signals while the anisotropic dielectric constant enhances the magnetic field in the c-axis (Mett et al., 2019), (ii) the oversized cavity shield minimizes the effects of the shield on dielectric resonator frequency and ohmic losses (Mett et al., 2008, 2019), (iii) the use of an oversized iris in conjunction with a Gordon coupler equipped with a Rexolite plug provides a purely resistive coupling mechanism that maintains frequency stability during adjustments (Mett & Hyde, 2022), and (iv) non-cylindrical sample tube geometries reduce the electric field within the aqueous sample, increasing the EPR signal acquired for the same sample volume (Sidabras et al., 2017).

The system demonstrates exceptional efficiency in sample utilization, requiring only ∼34 µL per shot. Spin-labeled protein concentrations as low as 12 µM were used in these experiments, establishing a new benchmark for sample economy in rapid-mixing EPR experiments. This advancement is particularly significant for studies involving samples that are difficult to produce in large quantities, where material conservation is paramount. Lastly, the experimentally determined dead time of 3.46 ms provides access to kinetics on nearly the entire ms timescale.

SF profiles of the TEMPOL-sodium dithionite reaction obtained at varying flow rates (Fig. 8A) collectively validate three critical aspects of system performance: the reproducibility of the mixing process, minimal mechanical vibration effects on signal quality, and the precise flow rate control provided by the driver system. The consistency of these measurements underscores the robust performance capabilities of our SF EPR system for kinetic measurements in this time regime. Incomplete mixing at slower flow rates is expected to generate non-linearity in the plot of steady-state signal amplitude vs. inverse flow rate. This phenomenon was not observed here, because the preset flow rate limitations of the μSFM are designed to maintain mixing efficiency (Fig. 3C); the results indicate that mixing efficiency is maintained even at the minimum tested flow rate of 0.7 mL/s.

### 3.2 Site-specific folding kinetics of T4L

To evaluate the capacity of the SF EPR system for large-scale protein studies, we investigated the folding kinetics of T4L at three distinct regions of the protein. A total of 198 individual relaxation profiles were collected and used to generate a chevron plot for each site, which were fit to a four-state model previously reported for the kinetic folding pathway of T4L (Cellitti, Bernstein, et al., 2007). The unfolding intermediate (J)—which generates curvature on the unfolding arm of the chevron plot—was previously reported to consist of a folded C-terminal subdomain and an unfolded N-terminal subdomain (Cellitti, Bernstein, et al., 2007).

The kinetic parameters obtained from site-directed spin labeling of T4L variants elucidate the folding dynamics of its subdomains, though substantial experimental uncertainties, reflected in the broad 95% confidence intervals in Table 1, warrant cautious interpretation. The C-terminal construct V131R1 displays a faster relative folding rate (*k*_*IJ*_ = 25.8 s^-1^), suggesting this subdomain drives the formation of an intermediate during early folding events, consistent with a previously proposed hierarchical model emphasizing a structured C-terminal intermediate (Cellitti, Bernstein, et al., 2007). In contrast, the N-terminal construct (S44R1) demonstrates the highest equilibrium stability (*K*_*UI*_ = 2.5 x 10^4^), indicating a significant role in the native state’s overall stability. However, the slower folding rate (*k*_*IJ*_= 8.9 s^-1^), when compared with V131R1, suggests that this region stabilizes the native state rather than initiating folding. Lastly, the equilibrium and rate constants derived from N68R1 generally reflect intermediate values between V131R1 and S44R1, coupling the N- and C-terminal subdomains.

All constructs exhibit extremely low equilibrium constants for the unfolding intermediate *K*_*NJ*_ ≈ 10^-8^, confirming the overwhelming stability of the native state under 0 M urea conditions. These results corroborate previous findings regarding the autonomous folding capability of the C-terminal region (Cellitti, Bernstein, et al., 2007; Cellitti, Llinas, et al., 2007; Kato et al., 2007; Llinás & Marqusee, 1998) while highlighting differences in subdomain-specific contributions to stability and dynamics. Although broad confidence intervals preclude definitive conclusions about unfolding kinetics, these data provide further evidence for a hierarchical folding mechanism involving intermediates on both sides of the rate-limiting transition state.

In some cases, the kinetic relaxation profiles exhibit a distinct bi-exponential decay upon refolding at lower urea concentrations, comprising an initial rapid exponential decay followed by a slower phase in which the signal amplitude increased. This biphasic behavior has been reported for the kinetic folding of multiple different proteins (Andersen et al., 2012; Garg et al., 2022; Kumar, 2016; Otzen & Andersen, 2013) and suggests distinct structural transitions during the folding process: an initial rapid phase corresponding to α-helical formation, followed by a slower phase associated with tertiary structure organization and stabilization. This feature is predominantly observed in the N68R1 construct, with similar features that are smaller in magnitude manifesting in the V131R1 and S44R1 constructs. This points to region-dependent variation in the T4L folding pathway and conformational dynamics not captured in the traditional chevron plot analysis.

These T4L folding results demonstrate the capability of the SF EPR system for elucidating local kinetic information and conservation of sample. Detection limits are fundamentally determined by the magnitude of signal changes and desired signal-to-noise ratios. Herein, there are examples of measurements using only 12 μM protein in 32 μL shots yielding a robust signal-to-noise ratio of 32.7. This technical achievement underscores the sensitivity of the system and efficiency in protein kinetics studies. The versatility of SF EPR detection extends beyond basic folding studies, offering potential applications in investigating local kinetic processes triggered by various biological factors, including denaturants, binding partners, lipids, membranes, and protein-protein interactions. What is particularly significant about the system is the capacity to operate with minimal sample quantities, making it valuable for studies involving proteins with limited availability or challenging expression profiles. To validate this capability, we extended our investigation to the more challenging β2AR, demonstrating the broad applicability of this methodology in protein dynamics research.

### 3.3 β2AR protein application

GPCRs are a large and diverse class of cell surface receptors responsible for regulating nearly every physiological process in the human body (Belmonte & Blaxall, 2012). While the solved structures from X-ray crystallography and cryo-electron microscopy provide important insight into the structural basis for GPCR signaling, growing recognition of the regulation of receptor signaling through modulation of the rates of the interconversion of conformational states (dynamics) (Du et al., 2019; García-Nafría & Tate, 2019; Gregorio et al., 2017; Liu et al., 2019; Manglik et al., 2015; Sandhu et al., 2019; Wingler & Lefkowitz, 2020)(Du et al., 2019; García-Nafría & Tate, 2019; Gregorio et al., 2017; Liu et al., 2019; Manglik et al., 2015; Sandhu et al., 2019; Wingler & Lefkowitz, 2020) underscores the importance of time-resolved measurements in defining the molecular mechanisms governing receptor activation.

As a proof of concept, we monitored the conformation of TM6 as a canonical marker for β2AR activation by the agonist BI-167107. The β2AR is a primary mediator of cardiopulmonary function and an important drug target for a variety of diseases including heart failure (Noor et al., 2011; Wang et al., 2018)(Noor et al., 2011; Wang et al., 2018) and asthma (Wendell et al., 2020)(Wendell et al., 2020). It is also a prototypical GPCR that serves as a useful model for many aspects of GPCR signaling and pharmacology. Given that the equilibrium exchange rate for the inward-outward equilibrium of TM6 is anticipated to be on the order of ms, the observed relaxation time constant of ∼260 s likely reflects rate of BI-167107 binding. This time-resolved measurement of an allosteric conformational change triggered by binding of an activating ligand illustrates the ability of the SF EPR system to access kinetic parameters in proteins that are challenging to express and analyze.

## 4. Conclusions

The high-sensitivity SF EPR system presented here enables investigation of protein conformational dynamics on the ms timescale with significantly reduced sample requirements compared with existing instruments. The system incorporates a custom dielectric resonator, optimized sample cell geometry, and an integrated SF mixer assembly with a CW EPR spectrometer to monitor time-resolved conformational changes with superior detection sensitivity. The utility of this system was demonstrated through two applications that highlight its versatility. A thorough analysis of T4L folding kinetics revealed previously unreported site-specific variation in the folding pathway. Additionally, successful measurement of ligand-induced conformational changes in the β2AR demonstrates the applicability to challenging membrane protein systems, where sample availability has traditionally been a significant barrier. Spin-label motion was used as a monitor of protein conformational equilibria in these examples, yet the use of CW EPR enables alternative detection modalities including solvent accessibility and distance measurements through dipolar or relaxation broadening. With the availability of this system, SF EPR can be readily extended to exploring conformational dynamics of complex, biomedically relevant proteins in various biological processes, including protein-protein and protein-ligand interactions.

## 5. Materials and Methods

### 5.1 T4L preparation

Site-directed mutagenesis of the T4L gene in the pET11a plasmid was performed using the QuikChange® site-directed mutagenesis kit (Agilent; Santa Clara, California) according to manufacturer protocols and verified by Sanger sequencing (Retrogen; San Diego, CA). T4L constructs used for EPR experiments include pseudo-wildtype (*WT) mutations C54T and C97A along with one of the following cysteine substitutions: V131C, N68C, or S44C.

T4L mutants (V131C, N68C, and S44C) were expressed and purified following established protocols (Langen et al., 2000; Sauers et al., 1992)(Langen et al., 2000; Sauers et al., 1992). Mutant plasmids were transformed into the expression cells *Escherichia coli* BL21(DE3) (Novagen; Madison, Wisconsin) and cultured under ampicillin-selective conditions at 37°C. Single colony transformants were grown overnight (16–18 h) in 15 mL Luria Broth containing 100 µg/mL ampicillin (LB-amp). The overnight culture was used to inoculate 1 L of LB-amp and grown to an OD600 of 0.8–1.0 over 3–4 h, followed by induction with 1 mM isopropyl-β-D-thio-galactopyranoside (Thermo Fisher Scientific; Waltham, Massachusetts) and expression for 1.5 h. Cells were harvested by centrifugation (6000 g, 20 min, 4°C), resuspended in buffer low-salt buffer (25 mM Tris, 25 mM MOPS, 0.1 mM EDTA, pH 7.6), and stored at −20°C until purification.

For protein purification, the thawed pellet was lysed by 3–4 passes through a French pressure cell at 900–1,000 psi. Soluble T4L was separated from cellular debris by centrifugation (36,600 g, 30 minutes, 4°C) and filtered through a 0.2 μm Whatman™ filter (Cytiva; Marlborough, Massachusetts). The clarified lysate was loaded onto a HiTrap SP HP cation exchange column equilibrated with low-salt buffer containing 5 mM dithiothreitol. T4L was eluted using a 1 M NaCl gradient, and the fractions corresponding to its molecular weight (as judged by SDS-PAGE) were pooled. T4L was either spin-labeled immediately after purification or stored in 20% glycerol (v/v) at −20°C for subsequent spin-labeling.

Dithiothreitol was removed immediately prior to spin labeling using a HiPrep 26/10 desalting column equilibrated with spin labeling buffer (50 mM MOPS, 25 mM NaCl, pH 6.8). Following desalting, purified T4L was spin-labeled with a 10-fold molar excess of MTSSL (2,2,5,5-tetramethyl-pyrroline-1-oxyl methanethiosulfonate) overnight at 4°C in the dark. Excess spin label was removed using a HiPrep 26/10 desalting column followed by five washes in a 10 kDa Ultra centrifugal filter (Amicon; Miami, Florida).

Spin-labeled T4L was concentrated to 100–150 µM as determined by A280 (ε = 24,750 cm⁻¹M⁻¹) measured via nanodrop (Thermo Fisher Scientific; Waltham, Massachusetts) and stored at 4°C for use within 24 h. Spin label concentration was determined using an ESR5000 CW EPR spectrometer (Bruker; Billerica, Massachusetts). Labeling efficiencies ranging from 0.6 to 1 spin per protein were observed for all T4L constructs. Protein purity exceeded 95% as assessed by SDS-PAGE.

For SF EPR experiments, T4L was pre-incubated in spin-label buffer with or without high concentrations of urea (see Tables S1–S3) to generate an initial equilibrium shifted strongly toward the natively folded (N) state or the unfolded (U) state, respectively.

### 5.2 β2AR preparation

Site-directed mutagenesis of the β2AR receptor gene in the pcDNA3.1-Zeo-tetO plasmid was performed using the QuikChange® site-directed mutagenesis kit (Agilent; Santa Clara, California) according to manufacturer protocols. All mutations were verified by Sanger sequencing (Retrogen; San Diego, California). The full length human β2AR construct used in these experiments contains the following mutations: five cysteine substitutions (C77V, C327S, C341L, C378A, and C406A) to avoid off-target labeling, two methionine substitutions (M96T, M98T) to enhance expression, and N-terminal FLAG and C-terminal histidine tags for purification. The native cysteine at residue 265 was spin-labeled for EPR experiments. Previous characterization has confirmed that this construct maintains wild-type ligand binding and G protein coupling properties (Fung et al., 2009; Gregorio et al., 2017; Manglik et al., 2015).

HEK293 cells were cultured in suspension at 37°C in a 300 mL volume to an optimal density of 3.0 × 10^6^ cells/mL with >98% viability. For the transient transfection, ∼1 μg/μL plasmid DNA prepared via Maxiprep kit (Qiagen; Hilden, Germany) and polyethylenimine were separately diluted in Opti-MEM (Thermo Fisher Scientific; Waltham, Massachusetts) at a DNA:polyethylenimine ratio of 1 μg:3.5 μL per mL of cell culture. After a 5-min incubation, the DNA and polyethylenimine solutions were combined and incubated for no more than 20 min at room temperature. The resulting transfection mixture, comprising ∼7% of the total culture volume, was added dropwise directly to the 300 mL cell culture. Protein expression was induced 24 h post-transfection using 0.4 mg/mL doxycycline, 0.2 mM alprenolol, and 0.5 M sodium butyrate, solubilized in Hank’s Balanced Salt Solution (Thermo Fisher Scientific; Waltham, Massachusetts). Cells were harvested by centrifugation (3,500 g) 48–60 h after induction, frozen in liquid nitrogen, and stored at −80°C.

Cell membranes were isolated through hypotonic lysis in buffer (20 mM Tris-HCl at pH 7.4, 2 mM EDTA, 10 mM MgCl2, 2.5 U/mL benzonase, 5.25 μM leupeptin, 1 mM benzamidine, and 5 μM alprenolol) stirring for 45 min at room temperature and 45 min at 4°C. Membranes were pelleted by centrifugation at 4000 g for 20 min to remove soluble cellular debris. The receptor was solubilized from the pelleted membranes in buffer containing 20 mM HEPES, 100 mM NaCl, 10mM MgCl^2^, 2.5 U/mL benzonase, 5 µM alprenolol, 5.25 µM leupeptin, 1 mM benzamidine, 0.05% (w/v) cholesteryl hemisuccinate (CHS), and 1% (w/v) DDM (dodecyl-β-maltoside) at 4°C. The solution was homogenized by 40 strokes in a Dounce homogenizer (Fisher; Plano, Texas) on ice, followed by stirring for 1 h at room temperature and then 1 h at 4°C. Insoluble material was removed via centrifugation at 40,000 g for 30 min. The soluble fraction containing receptor was filtered using a 0.45 µm pore size cellulous nitrate filter (Nalgene; Rochester, New York)). Next, the following reagents were added to the filtered receptor: 2 mM CaCl^2^, 5.25 µM leupeptin, 1 mM benzamidine, 5 µM alprenolol. The receptor was then loaded onto an M1 anti-FLAG immunoaffinity resin column, where it was washed with five column volumes of high salt buffer (20 mM HEPES pH 7.4, 2mM CaCl^2^, 500mM NaCl, 2 µM alp, 5.25 µM leupeptin, 1 mM benzamidine, 0.1% DDM, 0.005% CHS) to remove loosely bound hydrophilic molecules. This was followed by washing with five column volumes of a low-salt buffer (20 mM HEPES pH 7.4, 2mM CaCl^2^, 100mM NaCl, 2 µM alp, 5.25 µM leupeptin, 1 mM benzamidine, 0.1% DDM, 0.005% CHS).

The receptor was labeled on-column using low-salt buffer containing 100 μM MTSL via rotating overnight at 1 rpm at 4°C. Following labeling, the resin was washed with five column volumes of low-salt buffer containing 20 μM atenolol to remove bound alprenolol, followed by five column volumes of ligand-free low-salt buffer to yield ligand-free receptor.

The receptor was eluted using ligand-free low-salt buffer with 5mM EDTA and 0.2 mg/mL FLAG peptide. Eluted receptor was concentrated using a 100-kDa ultra centrifugal filter (Amicon; Miami, Florida) and subjected to size exclusion chromatography (Cytiva; Malborough, Massachusetts) using a Superdex 200 increase 10/300 column (Millipore Sigma; Burlington, Massachusetts). Monomeric fractions were pooled and concentrated using a 100-kDa ultra centrifugal filter (Amicon; Miami, Florida) to ∼50 µM as determined by A280 (ε = 65,235 cm⁻¹M⁻¹).

### 5.3 CW EPR spectroscopy

CW EPR spectra were recorded at X-band on an ESR5000 spectrometer (Bruker; Billerica, Massachusetts) at room temperature operating with the following parameters: 337 mT center field, 10 mT scan width, 41 s sweep time, 30 scans, 100 kHz modulation frequency, 0.1 mT modulation amplitude, 36.3 mW incident microwave power, and 9.46 GHz microwave frequency. For CW EPR measurements, 15 μL spin-labeled protein was loaded into a quartz capillary (0.9 mm ID × 1.4 mm OD). CW EPR data were area-normalized and plotted using custom Python software (available upon request from the authors).

### 5.4 SF EPR spectroscopy

Time-resolved EPR spectra were recorded at room temperature on an ELEXSYS 500 spectrometer (Bruker; Billerica, Massachusetts) using the SF EPR system described in the *Results* section. EPR signal intensity was monitored at a single magnetic field position using the following parameters: 10 mW incident microwave power, 100 kHz modulation frequency, 0.3 mT modulation amplitude, 1.28 ms time constant, and 9.33–9.45 GHz microwave frequency. Standard field-swept CW EPR spectra were collected of the reactants and products to determine optimal magnetic field position for monitoring the kinetics of each reaction.

For each set of experiments, the SF system was prepared according to a systematic cleaning and priming protocol. First, the two 500 µL Hamilton syringes are cleaned, loaded with the appropriate buffer, and mounted onto the driver system. The system was then primed with 34 µL of buffer from each syringe, with careful attention to eliminate any air bubbles. Following successful priming, reactants were loaded into the syringes, which were then secured with the pushing block precisely positioned against the syringe plunger.

Three parameters were programmed through BioLogic’s Bio-Kine software to define each SF experiment: shot volume, flow rate, and valve lead time. A total shot volume of 34 µL was used with a reactant mixing ratio of 1:1 to 1:9, depending on the desired final sample concentration and volumes being used. Flow rates varied from 0.7 to 1.8 mL/s. A valve lead time of 5 ms was used in all experiments to minimize the flow braking time while avoiding microphonic signal artifacts. A single shot was dispensed prior to data acquisition to verify the absence of air bubbles. The complete signal trace, encompassing all phases from initial flow to final decay, is recorded by the spectrometer’s data acquisition system for subsequent analysis.

### 5.5 System dead time evaluation using TEMPOL-dithionite reaction kinetics

To evaluate the system dead time, the center line minimum of a 1 mM TEMPOL spectrum was monitored over time following reduction with 10 mM sodium dithionite (Fig. 8). Analysis of the rate of TEMPOL reduction by sodium dithionite was done using a compressed exponential function, sometimes referred to as a Kohlrausch-Williams-Watts function,

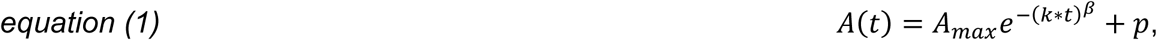

where *A*_*max*_ is the maximum signal at *t* = 0, *k* is the decay rate constant, *t* is time, *β* is the compressing factor, and *p* is the plateau. The fitted *k* value of 47.37 s^-1^ is consistent with previous reports for the reduction of TEMPOL and sodium dithionite using similar conditions.

In plots of the observed maximum signal vs. the inverse of the flow rate, *A*_*max*_ was determined by averaging the data points during the stationary phase of the reaction. The theoretical peak signal at infinite flow rate (*A*_0_) was determined to be 57 via a linear fit extrapolated to infinite flow rate (Fig. 3B). Using this *A*_0_ value and the *k* and *A*_*max*_ values obtained from our fastest flow rate of 1.8 mL/s, *equation 2* was used to extract a dead time of ∼3.46 ms:

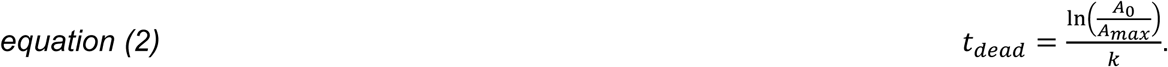

### 5.6 T4L SF EPR experimental design and data analysis

Most individual relaxation profiles were well-fit to a single exponential decay,

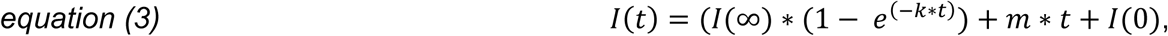

where *I*(*t*) is the EPR signal intensity at time t, *I*(0) represents the intensity at time zero, *I*(∞) is the intensity value asymptotically approached at infinite time, *k* is the relaxation rate constant, and *m* is the slope of the linear component attributed to baseline drift. Analysis of relaxation profiles was performed using custom Python scripts with the scipy.optimize library. The baseline drift was only significant when data collection time exceeded 15 min for a small subset of experiments.

In some cases, a bi-exponential decay was required to adequately fit the relaxation profiles,

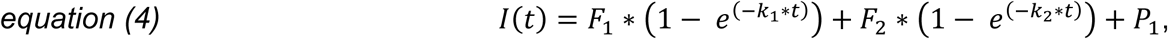

where *F*_1_ and *F*_2_ are the magnitude of the contribution to the first exponential term, *k*_1_ and *k*_2_ are the rate constants for their respective exponential component, and *P*_1_ and *P*_2_are the plateaus for each exponential component. The kinetic data revealed bi-exponential features predominantly in the lower concentrations of the folding arm. When time traces exhibited both fast and slow decay components across a range of urea concentrations, these components were grouped separately as they represented distinct kinetic processes. This systematic approach allowed us to track the transition from bi-exponential to single-exponential behavior with increasing denaturant concentration. In all cases, the slower rate was used in the chevron plot (Fig. S4). Importantly, bi-exponential fits did not benefit from including a linear term, as signal drift from the spectrometer was negligible at the time-scale of the observed relaxation profiles.

The complete dataset comprises both “folding” and “unfolding” experiments in which rapid mixing was used to either rapidly reduce or raise the urea concentration, respectively, and the relaxation to a new equilibrium was followed in real time by CW EPR as described previously. Plots of logarithmic folding and unfolding rates versus final urea concentration (i.e., chevron plots) were generated for each T4L construct and fit to a four-state sequential folding pathway as described below (Cellitti, Bernstein, et al., 2007; Cellitti, Llinas, et al., 2007; Llinás & Marqusee, 1998).

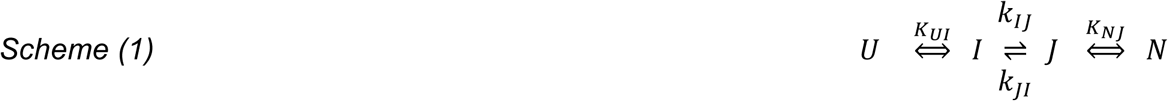

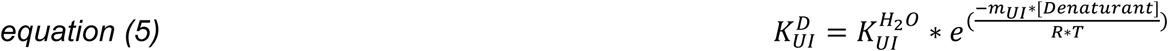

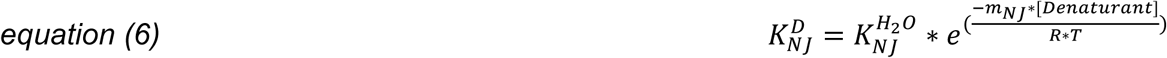

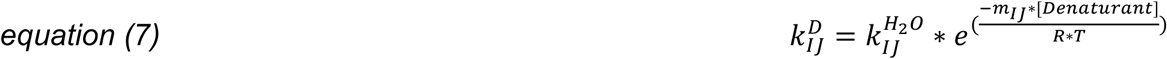

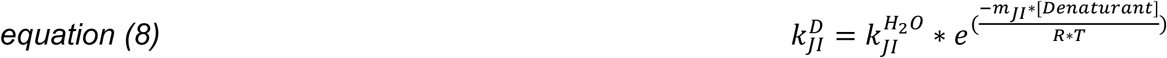

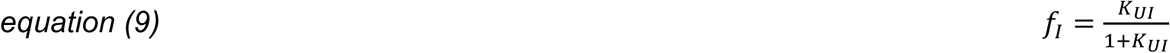

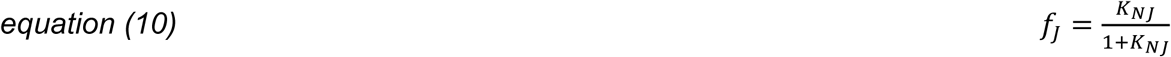

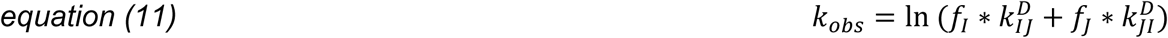

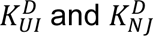 are the equilibrium constants for the U⇌I and N⇌J transitions, respectively. 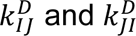 are the forward and reverse rate constants for the rate-limiting transition between intermediates I and J. *m* values represent the denaturant dependence of each transition, and [*Denaturant*] is the final concentration of urea after mixing. *f*_*I*_ and *f*_*J*_ represent the fractional populations of states I and J, respectively. Fits to the experimental data were performed using nonlinear least squares minimization and curve fitting in Python with the lmfit library. Details of all experimental conditions— including urea and protein concentrations, injection volumes, flow rates, and dead times—are provided in Supplemental Tables S1–S3.

### 5.7 β2AR SF EPR experimental design and data analysis

Solutions of 50 µM β2AR C265R1 and 50 µM BI-167107 were prepared in identical buffers [20 mM HEPES, 100 mM NaCl, 0.05% (w/v) CHS, 1% (w/v) DDM]. The reactants were mixed at a 1:1 ratio at a flow rate of 1.8 mL/s, and the CW EPR signal intensity at the minimum of the center field line was monitored in real time. The relaxation profile was well-fit to a single exponential decay (Equation 3).

## Supplementary Material

The Supplementary Material includes experimental parameters for stopped-flow electron paramagnetic resonance (EPR) studies conducted on three T4 Lysozyme (T4L) constructs (S44R1, N68R1, V131R1). These tables detail reaction conditions, instrumental configurations, and kinetic acquisition settings (tables S1-S3). The mean of the natural logarithm of the rate constants (ln(*k*)) for each T4L construct, accompanied by their corresponding 95% confidence intervals is provided in tables S4-S6. Figure S1 shows the stopped-flow EPR time-resolved data illustrating the reaction kinetics of 1 mM TEMPOL with 50 mM sodium dithionite across flow rates of 0.7–1.8 mL/s. The equilibrium peak signal amplitude is shown to depend on the flow rate, reflecting mixing efficiency and reaction completion. Figure S2 displays X-band continuous-wave EPR (CW-EPR) spectra of T4L constructs S44R1, N68R1, and V131R1 under equilibrium conditions in the absence (0 M) and presence (9 M) of urea. These spectra highlight urea-induced conformational changes in each construct. Figure S3, shows the Mathematical models (equations) applied to fit the kinetic traces obtained from the reaction of 1 mM TEMPOL with 50 mM sodium dithionite at a flow rate of 0.7 mL/s.

(SF_EPR_Supplementary_Material.pdf

## Acknowledgements

Research reported in this publication was supported by the National Institute of General Medical Sciences of the National Institutes of Health under award number GM140385 (CSK and MTL). The content is solely the responsibility of the authors and does not necessarily represent the official views of the National Institutes of Health. The authors thank Timothy Thelaner and Richard Scherr for technical expertise in machining and constructing the resonator and mixer components. We are grateful for the help of Michael D. Bridges, PhD, in defining certain equations used in the T4L kinetic analysis. We are also grateful for the help of Michael Vaughn (PhD) for his willingness to answer any questions we may have had regarding the Biologic (Seyssinet-Pariset, France) μSFM system.

## Author Contributions

Alexander M. Garces: Writing – original draft; writing – review and editing; investigation; methodology; data curation; formal analysis; software; visualization.

Richard R. Mett: Methodology; resources; funding acquisition.

Candice S. Klug: Writing – review and editing; conceptualization; resources; funding acquisition; supervision.

Jason W. Sidabras: Writing – original draft; writing – review and editing; methodology; visualization; resources; funding acquisition.

Michael T. Lerch: Writing – original draft; writing – review and editing; conceptualization; validation; resources; funding acquisition; supervision.

## Conflict of Interest Statement

The authors declare no conflicts of interest.

## Notes

### Competing Interest Statement

The authors have declared no competing interest.

