## Supplementary Material for "A High-Sensitivity Stopped-Flow EPR System to Monitor Millisecond Conformational Kinetics in Spin-Labeled Proteins"

**Table S1: T4L S44R1 Experimental Parameters**

| Final Urea Conc. (M) | Start Urea w/T4L (M) | Start Urea w/o T4L (M) | Start Protein Conc. (μM) | Protein Vol (μl) | Buffer Vol (μl) | Flow (ml/s) | Shot Time (ms) | Dead Time (ms) | Rep. |
|---|---|---|---|---|---|---|---|---|---|
| 2.20 | 8 | 1 | 150 | 6 | 29 | 1.1 | 31.82 | 6.29 | 2 |
| 2.65 | 9 | 0 | 150 | 10 | 24 | 1.2 | 28.33 | 5.77 | 3 |
| 3.06 | 8 | 1 | 150 | 10 | 24 | 1.2 | 28.33 | 5.77 | 3 |
| 3.44 | 9 | 0 | 150 | 13 | 21 | 1.4 | 24.29 | 4.94 | 4 |
| 3.97 | 9 | 0 | 150 | 15 | 19 | 1.6 | 21.25 | 4.33 | 4 |
| 3.91 | 1 | 9 | 150 | 21 | 12 | 1.4 | 23.57 | 4.94 | 3 |
| 4.00 | 8 | 1 | 150 | 15 | 20 | 1.5 | 23.33 | 4.61 | 3 |
| 4.50 | 9 | 0 | 150 | 17 | 17 | 1.8 | 18.89 | 3.84 | 7 |
| 5.00 | 1 | 9 | 150 | 17 | 17 | 1.8 | 18.89 | 3.84 | 3 |
| 5.47 | 9 | 1 | 150 | 19 | 15 | 1.6 | 21.25 | 4.33 | 5 |
| 5.00 | 8 | 1 | 150 | 20 | 15 | 1.5 | 23.33 | 4.61 | 4 |
| 5.47 | 1 | 9 | 150 | 15 | 19 | 1.6 | 21.25 | 4.33 | 4 |
| 5.94 | 1 | 9 | 150 | 13 | 21 | 1.4 | 24.29 | 4.94 | 3 |
| 6.00 | 8 | 1 | 150 | 25 | 10 | 1.2 | 29.17 | 5.77 | 2 |
| 6.65 | 1 | 9 | 150 | 10 | 24 | 1.2 | 28.33 | 5.77 | 3 |
| 7.06 | 1 | 9 | 150 | 8 | 25 | 1.2 | 27.50 | 5.77 | 5 |
| 7.49 | 1 | 9 | 150 | 7 | 30 | 1.1 | 33.64 | 6.29 | 3 |
| 7.94 | 0 | 10 | 150 | 7 | 27 | 1.1 | 30.91 | 6.29 | 5 |
| 8.53 | 0 | 10 | 150 | 5 | 29 | 1.0 | 34.00 | 6.92 | 5 |
| 8.75 | 0 | 10 | 150 | 4 | 28 | 1.0 | 32.00 | 6.92 | 4 |

**Table S2: T4L N68R1 Experimental Parameters**

| Final Urea Conc. (M) | Start Urea w/T4L (M) | Start Urea w/o T4L (M) | Start Protein Conc. (μM) | Protein Vol (μl) | Buffer Vol (μl) | Flow (ml/s) | Shot Time (ms) | Dead Time (ms) | Rep. |
|---|---|---|---|---|---|---|---|---|---|
| 1.94 | 6 | 0 | 150 | 11 | 23 | 1.3 | 26.15 | 5.32 | 3 |
| 2.16 | 8 | 0 | 150 | 10 | 27 | 1.2 | 30.83 | 5.77 | 3 |
| 2.47 | 6 | 0 | 150 | 14 | 20 | 1.5 | 22.67 | 4.61 | 3 |
| 2.47 | 8 | 1 | 150 | 8 | 30 | 1.1 | 34.55 | 6.29 | 3 |
| 3.06 | 8 | 0 | 150 | 13 | 21 | 1.4 | 24.29 | 4.94 | 2 |
| 3.06 | 8 | 1 | 150 | 10 | 24 | 1.2 | 28.33 | 5.77 | 3 |
| 3.53 | 8 | 0 | 150 | 15 | 19 | 1.6 | 21.25 | 4.33 | 3 |
| 3.59 | 9 | 1 | 100 | 11 | 23 | 1.3 | 26.15 | 5.32 | 3 |
| 4.06 | 9 | 1 | 100 | 13 | 21 | 1.4 | 24.29 | 4.94 | 4 |
| 4.53 | 9 | 1 | 100 | 15 | 19 | 1.6 | 21.25 | 4.33 | 3 |
| 5.00 | 1 | 9 | 100 | 17 | 17 | 1.8 | 18.89 | 3.84 | 2 |
| 5.00 | 9 | 1 | 100 | 17 | 17 | 1.8 | 18.89 | 3.84 | 2 |
| 5.47 | 9 | 1 | 100 | 19 | 15 | 1.6 | 21.25 | 4.33 | 3 |
| 5.47 | 1 | 9 | 100 | 15 | 19 | 1.6 | 21.25 | 4.33 | 2 |
| 5.94 | 1 | 9 | 100 | 13 | 21 | 1.4 | 24.29 | 4.94 | 3 |
| 5.94 | 9 | 1 | 100 | 21 | 13 | 1.4 | 24.29 | 4.94 | 5 |
| 6.58 | 1 | 9 | 100 | 10 | 23 | 1.2 | 27.50 | 5.77 | 3 |
| 7.06 | 0 | 10 | 100 | 10 | 24 | 1.2 | 28.33 | 5.77 | 5 |
| 7.58 | 0 | 10 | 100 | 8 | 25 | 1.2 | 27.50 | 5.77 | 3 |
| 7.94 | 0 | 10 | 100 | 7 | 27 | 1.1 | 30.91 | 6.29 | 4 |
| 8.53 | 0 | 10 | 100 | 5 | 29 | 1.0 | 34.00 | 6.92 | 3 |
| 8.82 | 0 | 10 | 100 | 4 | 30 | 1.0 | 34.00 | 6.92 | 3 |

**Table S3: T4L V131R1 Experimental Parameters**

| Final Urea Conc. (M) | Start Urea w/T4L (M) | Start Urea w/o T4L (M) | Start Protein Conc. (μM) | Protein Vol (μl) | Buffer Vol (μl) | Flow (ml/s) | Shot Time (ms) | Dead Time (ms) | Rep. |
|---|---|---|---|---|---|---|---|---|---|
| 1.94 | 8 | 0 | 150 | 8 | 25 | 1.1 | 30 | 6.29 | 2 |
| 2.44 | 8 | 1 | 150 | 7 | 27 | 1.1 | 30.91 | 6.29 | 2 |
| 3.06 | 8 | 1 | 150 | 10 | 24 | 1.2 | 28.33 | 5.77 | 2 |
| 3.59 | 9 | 1 | 150 | 11 | 23 | 1.3 | 26.15 | 5.32 | 3 |
| 4.06 | 9 | 1 | 150 | 13 | 21 | 1.4 | 24.29 | 4.94 | 6 |
| 4.53 | 9 | 1 | 150 | 15 | 19 | 1.6 | 21.25 | 4.33 | 3 |
| 5 | 9 | 1 | 150 | 17 | 17 | 1.8 | 18.89 | 3.84 | 6 |
| 5.47 | 9 | 1 | 150 | 19 | 15 | 1.6 | 21.25 | 4.33 | 6 |
| 5.94 | 9 | 1 | 150 | 21 | 13 | 1.6 | 21.25 | 4.33 | 3 |
| 5.94 | 1 | 9 | 150 | 13 | 21 | 1.4 | 24.29 | 4.94 | 2 |
| 6.58 | 1 | 9 | 150 | 10 | 23 | 1.2 | 27.50 | 5.77 | 3 |
| 7.06 | 0 | 10 | 100 | 10 | 24 | 1.1 | 30.91 | 6.29 | 1 |
| 7.06 | 0 | 10 | 100 | 10 | 24 | 1.2 | 28.33 | 5.77 | 2 |
| 7.06 | 1 | 9 | 150 | 8 | 25 | 1.1 | 30 | 6.29 | 2 |
| 7.58 | 0 | 10 | 100 | 8 | 25 | 1.2 | 27.50 | 5.77 | 2 |
| 7.94 | 0 | 10 | 100 | 7 | 27 | 1.1 | 30.91 | 6.29 | 2 |
| 7.94 | 0 | 10 | 100 | 7 | 27 | 1 | 34 | 6.92 | 4 |
| 8.53 | 0 | 10 | 100 | 5 | 29 | 1 | 34 | 6.92 | 1 |
| 8.82 | 0 | 10 | 100 | 4 | 30 | 1 | 34 | 6.92 | 3 |

**Table S4: T4L S44R1 Rate Constants for Kinetic Traces**

| [Urea] (M) | ln($k_1$) | ln($k_2$) |
|---|---|---|
| 2.200 | 0.703 ± 0.824 | 1.999 ± 6.664 |
| 2.640 | 0.670 ± 0.598 | 1.742 ± 13.322 |
| 3.050 | 0.442 ± 0.087 | -- |
| 3.440 | 0.156 ± 0.035 | -- |
| 3.910 | -0.520 ± 0.033 | -- |
| 3.970 | -0.519 ± 0.003 | -- |
| 4.000 | -0.800 ± 0.001 | -- |
| 4.500 | -1.650 ± 0.000 | -- |
| 5.000 | -2.130 ± 0.000 | -- |
| 5.470 | -2.538 ± 0.000 | -- |
| 5.940 | -2.766 ± 0.000 | -- |
| 6.000 | -2.730 ± 0.000 | -- |
| 6.640 | -2.264 ± 0.000 | -- |
| 7.060 | -1.913 ± 0.000 | -- |
| 7.480 | -1.716 ± 0.000 | -- |
| 7.940 | -1.510 ± 0.002 | -- |
| 8.520 | -1.384 ± 0.001 | -- |
| 8.750 | -1.367 ± 0.001 | -- |

*Note: Values represent log of mean rate constants ± mean confidence interval for each urea concentration. Missing values (--) indicate no $k_2$ data were available for those concentrations.*

**Table S5: T4L N68R1 Rate Constants for Kinetic Traces**

| [Urea] (M) | ln($k_1$) | ln($k_2$) |
|---|---|---|
| 1.940 | 1.49506 ± 0.12179 | 27.004 ± 10.183 |
| 2.160 | 1.61550 ± 0.22423 | 15.924 ± 6.314 |
| 2.470 | 1.16525 ± 0.16117 | 8.884 ± 1.903 |
| 3.050 | 0.91168 ± 0.09720 | 6.527 ± 0.498 |
| 3.530 | 0.80078 ± 0.87420 | 3.882 ± 0.174 |
| 3.580 | 0.22551 ± 0.07540 | 6.235 ± 0.336 |
| 4.050 | 0.10527 ± 0.00063 | 4.522 ± 0.120 |
| 4.520 | 0.04431 ± 0.00007 | 5.057 ± 0.525 |
| 5.000 | 7.674 × 10^-3 ± 0.00003 | -- |
| 5.470 | 2.690 × 10^-3 ± 0.00001 | -- |
| 5.940 | 1.464 × 10^-3 ± 0.00001 | -- |
| 6.570 | 3.208 × 10^-3 ± 0.00001 | -- |
| 7.050 | 0.01272 ± 0.00004 | -- |
| 7.570 | 0.02127 ± 0.00021 | -- |
| 7.940 | 0.01917 ± 0.00014 | -- |
| 8.520 | 0.03930 ± 0.00048 | -- |
| 8.820 | 0.03888 ± 0.00049 | -- |

*Note: Values represent mean rate constants ± mean confidence interval for each urea concentration. Missing values (--) indicate no $k_2$ data were available for those concentrations.*

**Table S6: T4L V131R1 Rate Constants for Kinetic Traces**

| [Urea] (M) | $\ln(k_1)$ | $\ln(k_2)$ |
|---|---|---|
| 1.940 | 33.359 ± 12.527 | 4.89201 ± 1.48270 |
| 2.440 | 11.11791 ± 1.97972 | -- |
| 3.060 | 5.43497 ± 0.48191 | -- |
| 3.590 | 1.93050 ± 0.10129 | -- |
| 4.060 | 0.28652 ± 0.00332 | -- |
| 4.530 | 0.06519 ± 0.00023 | -- |
| 5.000 | 0.01852 ± 0.00004 | -- |
| 5.470 | 4.561 × 10^-3 ± 0.00001 | -- |
| 5.940 | 2.630 × 10^-3 ± 0.00008 | -- |
| 6.580 | 8.020 × 10^-3 ± 0.00001 | -- |
| 7.060 | 0.01054 ± 0.00003 | -- |
| 7.580 | 0.01407 ± 0.00007 | -- |
| 7.940 | 0.02213 ± 0.00013 | -- |
| 8.530 | 0.03263 ± 0.00035 | -- |
| 8.820 | 0.04130 ± 0.00044 | -- |

*Note: Values represent mean rate constants ± mean confidence interval for each urea concentration. Missing values (--) indicate no $k_2$ data were available for those concentrations.*

Figure S1 title: Complete SF EPR traces of 1 mM TEMPOL reduced by 50 mM sodium diothionite at flow rates from 0.7 to 1.8 mL/s. Traces are color-coded as indicated.

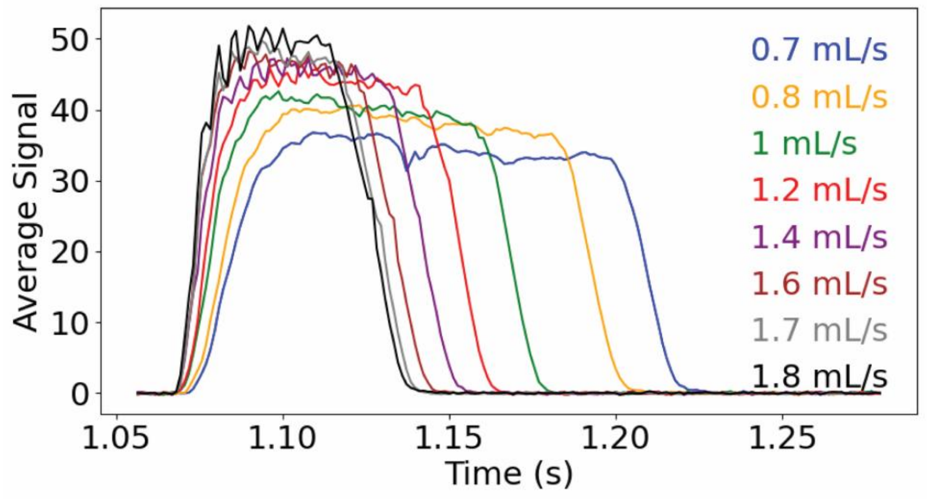


**Figure S1.** SF EPR data. Time-resolved data monitoring the peak of the low field line of the nitroxide spectrum during the reaction between 1 mM TEMPOL and 50 mM sodium dithionite at pH 7 at different flow rates ranging from 0.7 mL/s to 1.8 mL/s.

Figure S2 title: CW EPR spectra of T4L S44R1 (top), N68R1 (middle), and V131R1 (bottom) in 0 M and 9 M urea. Spectra are area-normalized.

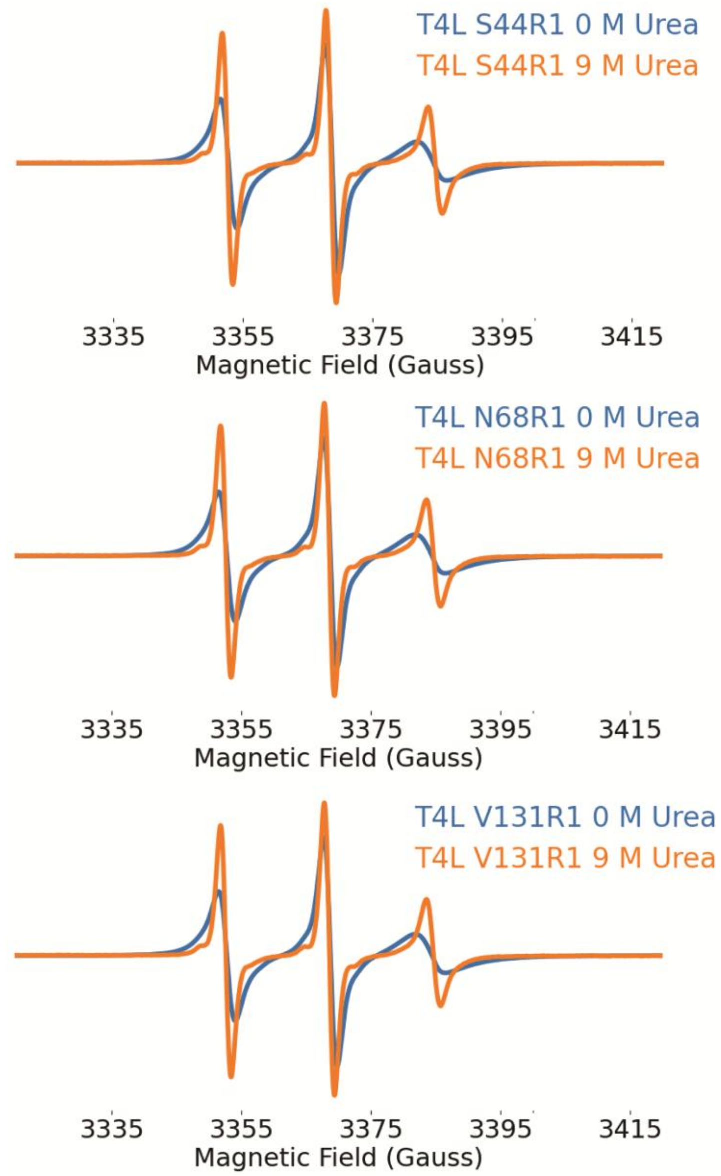


**Figure S2.** X-band CW-EPR collected at room temperature over 100 G spectra of T4L constructs S44R1, N68R1, and V131R1.

Figure S3 title: Evaluation of different models for fitting the kinetics of 1 mM TEMPOL reduction by 50 mM sodium dithionite at a flow rate of 0.7 mL/s. The signal decay after flow termination was fit using a stretched exponential (top), single exponential (middle), and linear (bottom) model.

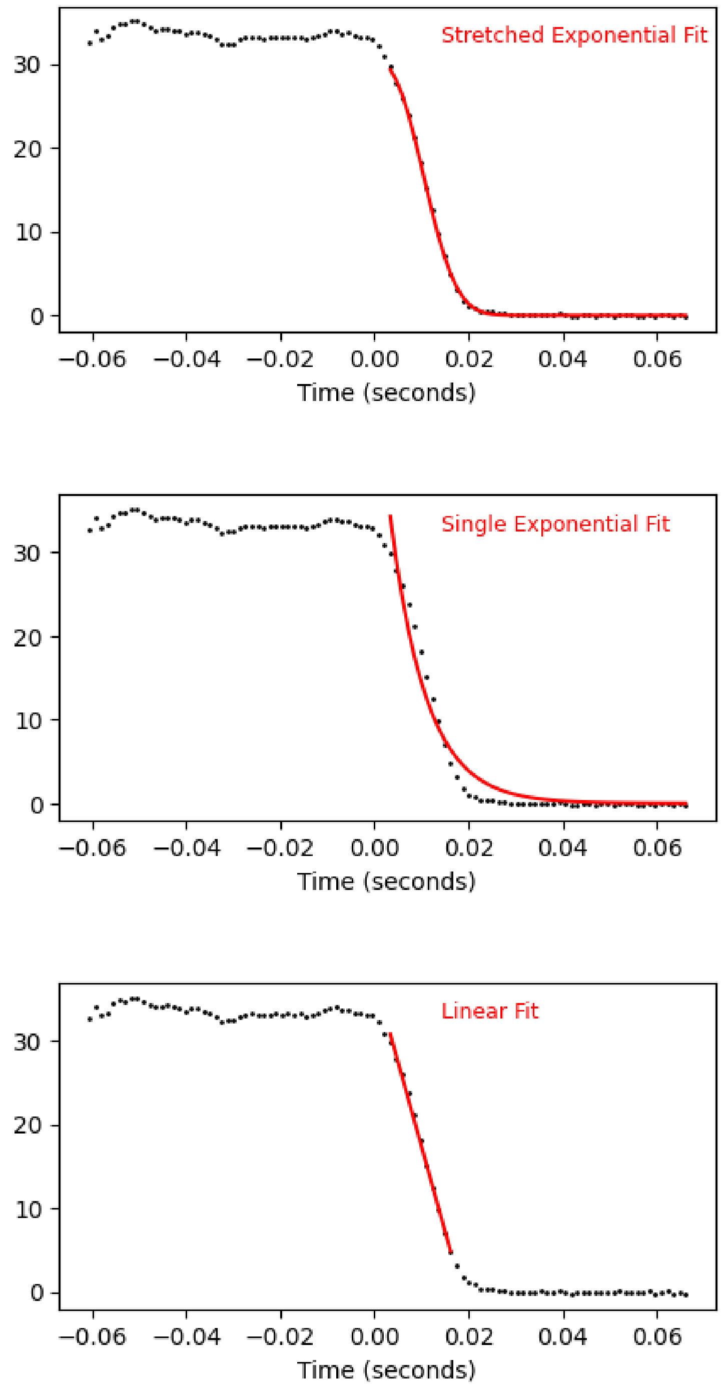


**Figure S3**. SF EPR spectrum of 1:1 mixing ratio of 1 mM TEMPOL reduced by 50 mM sodium dithionite at a flow rate of 0.7 mL/s.

Figure S4 title: Urea-dependent kinetics of T4L folding. The natural logarithm of the relaxation time constants determined from fits to individual SF EPR relaxation profiles are plotted versus urea concentration for (A) S44R1, (B) N68R1, and (C) V131R1. In cases where a bi-exponential fit was used, both the fast (red) and slow (blue) time constants are shown.

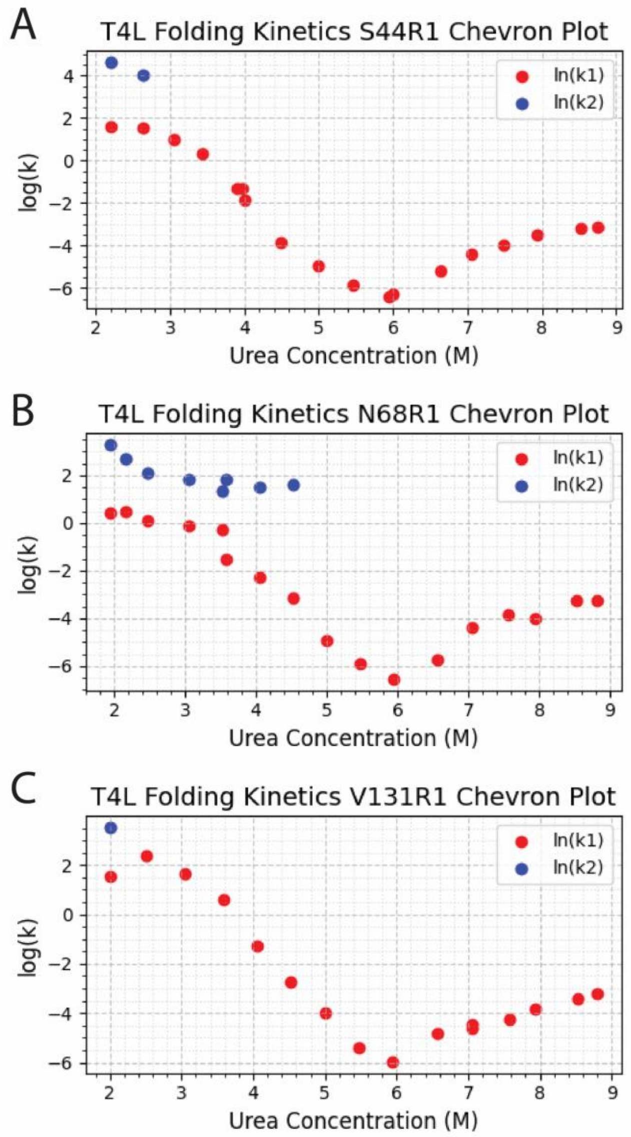


**Figure S4.** Chevron plots describing the kinetics of T4L folding at three distinct regions of the protein: H helix (V131R1), C helix (N68R1), and B helix (S44R1). k1 is shown in red and k2 is shown in blue. (**A**) Chevron plot of S44R1. (**B**) Chevron plot of N68R1. (**C**) Chevron plot of V131R1.
